# High-grade serous ovarian carcinoma organoids as models of chromosomal instability

**DOI:** 10.1101/2022.09.01.506155

**Authors:** Maria Vias, Lena Morrill Gavarró, Carolin M. Sauer, Debbie Sanders, Anna M. Piskorz, Dominique-Laurent Couturier, Stéphane Ballereau, Barbara Hernando, James Hall, Filipe Correia-Martins, Florian Markowetz, Geoff Macintyre, James D. Brenton

**Affiliations:** Cancer Research UK Cambridge Institute, University of Cambridge, Li Ka Shing Centre, Robinson Way, Cambridge CB2 0RE, UK; Centro Nacional de Investigaciones Oncológicas, C/Melchor Fernández Almagro, 3. 28029 Madrid

**Author notes:** Equal contribution (Maria Vias, Lena Morrill Gavarró).

## Abstract

High-grade serous ovarian carcinoma (HGSOC) is the most genomically complex cancer, characterised by ubiquitous *TP53* mutation, profound chromosomal instability and heterogeneity. The mutational processes driving chromosomal instability in HGSOC can be distinguished by specific copy number signatures. To develop clinically relevant models of these mutational processes we derived 15 continuous HGSOC patient-derived organoids (PDOs). We carried out detailed bulk transcriptomic, bulk genomic, single cell genomic, and drug sensitivity characterisation of the organoids. We show that PDOs comprise communities of different clonal populations and represent models of different causes of chromosomal instability including homologous recombination deficiency, chromothripsis, tandem-duplicator phenotype and whole genome duplication. We also show that these PDOs can be used as exploratory tools to study transcriptional effects of copy number alterations as well as compound-sensitivity tests. In summary, HGSOC PDO cultures provide a genomic tool for studies of specific mutational processes and precision therapeutics.

## Introduction

HGSOC is a heterogeneous, chromosomally unstable cancer with predominant somatic copy number alterations (SCNAs) and other structural variants including large-scale chromosomal rearrangements. Oncogenic mutations are rare and recurrent somatic substitutions involve less than 10 driver genes^1–4^. We have previously shown that copy-number signatures are able to recapitulate the major defining elements of HGSOC genomes and illuminate a fundamental structure underlying genomic complexity across chromosomally unstable human cancers^56^.

Improving outcomes in HGSOC will depend on having well-characterised and validated pre-clinical *in vitro* models, but currently available 2D models have multiple shortcomings such as changes in cell morphology, loss of diverse genotype and polarity, as well as other limitations. Patient-derived organoids (PDOs) offer improved pre-clinical cancer models and are generally are molecularly representative of the donor, have good clinical annotation and can represent tumoural intra-heterogeneity^7–10^. PDOs can be cultured for short periods^11,12^ but continuous HGSOC PDOs have only been generated for 27 models^10,13^ and these models lack detailed genomic characterization to determine whether they represent the broad landscape of genomic instability in HGSOC.

Approximately 50% of HGSOC patients may have impaired homologous-recombination (HR) DNA repair, including approximately 15% of cases that have loss of function and epigenetic events in *BRCA1* and *BRCA2*^2^. Consequently, homologous-recombination deficiency (HRD) is the major genomic classifier in the clinic and stratifies patients for outcome after treatment with PARP inhibitors^14,15^. Despite the relatively high prevalence of HRD and BRCA1/2 mutations in the clinic, there are only very few relevant models suggesting selection against cell lines and PDO that carry *BRCA1* and *BRCA2* deleterious mutations. In addition, there is an unmet clinical need for therapies in HGSOC that are homologous recombination proficient (HRP). Several distinctive patterns of structural variation have been described in HRP tumours including chromothripsis, tandem duplication (TD), whole-genome duplication (WGD) and *CCNE1* amplification^4^. Apart from the description of *CCNE1* amplification, it is unknown if the HGSOC organoids described to date display any of these genomic features and most cell line publications only refer to *BRCA1* and *BRCA2* mutations. These shortcomings highlight the lack of a systematic approach to characterize CIN and copy number signatures in PDO models.

To address these challenges, we developed HGSOC PDOs and characterised their genomes, transcriptomes, drug sensitivity and intra-tumoural heterogeneity. Using copy number signatures, we show that our models comprehensively recapitulate clinically relevant genomic features across the whole spectrum of CIN observed in HGSOC patients. PDOs showed strong copy-number-driven gene expression and transcriptional heterogeneity between models. Drug sensitivity was reproducible compared to parental tissues and the ability of these models to grow *in vivo*. Single cell DNA sequencing showed copy number features at a subclonal level and distinct clonal populations. The PDO models we present thus shed light on evolutionary characteristics of HGSOC and can have clinical relevance for guiding treatment decisions.

## Results

### HGSOC organoid culture derivation

To establish HGSOC organoids we used cells obtained from patient-derived ascites (n = 57), solid tumours (n = 14) and patient derived xenografts (n = 15) (Fig. 1a). Most ascites cultures were derived from patients with recurrent HGSOC and clinical summaries are provided in Supplementary Fig. 2 and Supplementary Table 1. We tested the effect of two published^10,16^ media compositions on 15 independent cultures and found similar PDO viability (Fig. 1b). We therefore performed subsequent derivations using the less complex fallopian tube media^16^. The efficiency of establishing PDOs was dependent on the type of tissue sample used for derivation (p<0.0001, log-rank test) and the highest success rate for short-term cultures (passage number between 1 and 4) was obtained using ascites and dissociated xenograft tissues (65%) (N = 86; Supplementary Fig. 1c). We defined continuous PDO cultures as those that could be serially passaged >5 times followed by cryopreservation and successful re-culture. Using these criteria, PDO were established for 15/18 organoid lines (PDO16, 17 and 18 were finite culture models). Four PDOs were able to grow as continuous 2D cell lines in conventional tissue culture media (Supplementary Table 2).

**Fig. 1.**
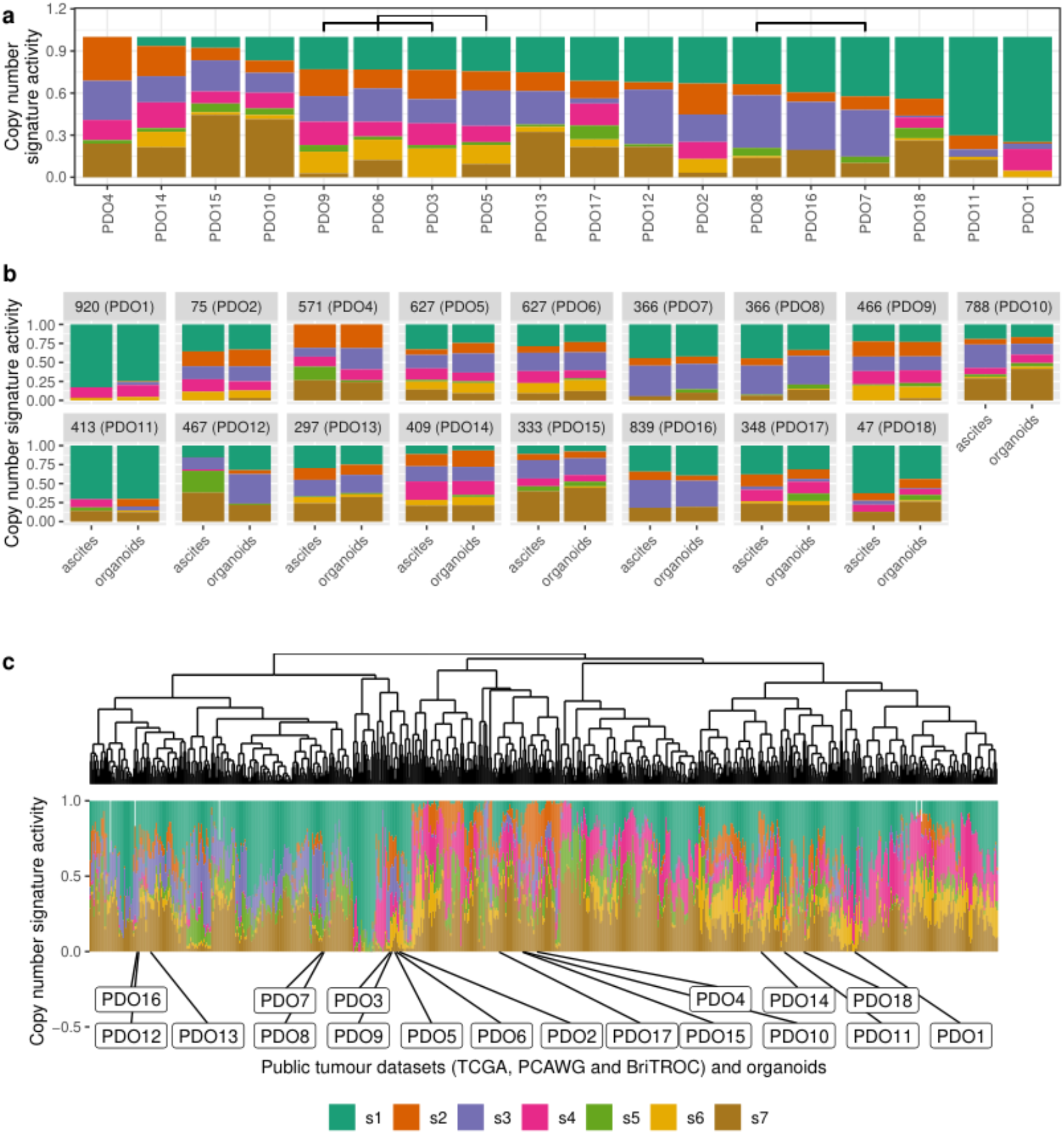
Chromosomal instability features of patient-derived organoids (PDOs) **a** Stacked bar plots show copy number signature activities ranked by signature s1 (PDO 16, 17 and 18 were not continuous models). Brackets indicate PDOs derived from the same individual. **b** Stacked bar plots show copy number signature activities for organoids and the matched ascites tissue sample from which they were derived **c** Unsupervised hierarchical clustering of copy number signature for PDO and 692 HGSOC cases using Aitchinson’s distance with complete linkage function. Stacked barplots in lower panel show copy number signature activities.

PDOs were screened for mutations enriched in HGSOC using an in-house tagged amplicon sequencing panel (Supplementary Fig. 3 and Supplementary Table 3) and were highly comparable to mutational profiles and p53 immunostaining from the original patient sample (Supplementary Fig. 4). Pathogenic somatic *BRCA1* or *BRCA2* mutations were present in PDO4, PDO7, PDO8 and PDO9 (Supplementary Fig. 3b and Supplementary Table 3). Germline DNA sequencing for 11 of the PDO donors (Supplementary Table 4) showed *BRCA1/2* germline mutations with unknown clinical significance or benign variants in patients OV04-297 (PDO13), OV04-409 (PDO14) and OV04-627 (PDO5 and PDO6).

To assess feasibility of the PDOs for *in vivo* modelling, we implanted 8 PDO models into immunodeficient mice using intraperitoneal injection to simulate peritoneal metastasis. All 8 PDOs efficiently established PDX models and 7/8 resulted in solid implants on peritoneal surfaces and/or liver infiltration (Supplementary Fig. 5).

### Genomic characterization of patient-derived organoids

We characterised the genomic landscape of the PDOs using shallow whole genome sequencing (sWGS) and derived copy number signatures to characterize the diversity of causes of CIN (Fig. 1a and Supplementary Fig. 6). We used seven previously identified copy number signatures in ovarian cancer that represent different putative causes of CIN: s1: mitotic errors, s2: replication stress causing tandem duplication, s3 and s7: homologous recombination deficiency, s5: unknown etiology leading to chromothripsis, and s6: replication stress leading to focal amplification. The finite lines PDO16, PDO17 and PDO18 are included here for comparison only.

PDO1 and PDO11 showed high levels of signature s1 and are thus appropriate models of mitotic errors. PDO4 exhibited high activity of a signature of replication stress induced tandem duplication (s2) but did not have a canonical *CDK12* mutation suggesting this may represent an alternative model of tandem duplication (see also below)^17,18^. Thirteen of the organoids showed evidence of s3 and can be considered as having HRD. Of these, pathogenic somatic *BRCA1* and *BRCA2* mutations were present in PDO4, PDO7, PDO8 and PDO9 (Supplementary Fig. 3b and Supplementary Table 3); a novel non-synonymous secondary mutation was observed in *BRCA1* (c.1367T>C) in PDO8 which was cultured after progression on PARP inhibitor therapy (paired with PDO7; Supplementary Fig. 2 and Supplementary Fig. 3); *BRCA1/2* mutations were not detected in the remaining PDOs with s3 (PDO2, PDO3, PDO10, PDO12, PDO15) suggesting these may be models of other mechanisms of HRD. PDO1 and PDO11 showed low signature s3 activity making them suitable models of HRP ovarian cancer. Ten of the PDOs showed s4 activity making them suitable to study the effects of WGD. Signature s5, with unknown etiology that results in chromothripsis had generally low activity in all PDOs consistent with previous observations suggesting chromothripsis is a rare event in HGSOC. s6, a signature of replication stress resulting in focal amplification was high in PDO3, PDO5, PDO6, PDO9 and PDO14 indicating these are good models to study both the cause and consequence of focal amplification events. Finally, a number of organoids showed s7 making them good models to study the effects of HRD following WGD.

### Organoids represent the spectrum of human high-grade serous ovarian cancers

We next compared copy number signatures from donor patient tissues and matched PDO (Fig. 1) and found that they were highly consistent except for PDO12 (OV04-467). For PDO12, the parental *CDK12* mutation present in the ascites specimen was not recovered after culture, suggesting selection for a subclonal population with distinct copy number signatures (Supplementary Fig. 3a).

Both PDO culture and derivation of PDX models may negatively select against specific molecular subtypes of HGSOC—which may explain the low number of *BRCA1/2* models. To test whether the PDOs were representative of the wider population of HGSOC cases, we compared PDO copy number features to those of publicly available patient cohorts (N = 692 samples from the TCGA, PCAWG and BriTROC-1 studies). The number of copy number segments (Supplementary Fig. 7a) did not significantly differ between PDOs (169 ± 77) and HGSOC tissues from TCGA, PCAWG and BriTROC-1 (200 ± 134) (p=0.22, negative binomial likelihood ratio test). Ploidy was found to be bimodal in both groups, with centers at average ploidies 2 and 3.5 (Supplementary Fig. 7b). There were also no significant differences in other copy number features (Supplementary Fig. 7c).

We next clustered copy number activity profiles (Fig. 1d) from TCGA, PCAWG and BriTROC (N = 692) and compared these with the PDO profiles. Unsupervised hierarchical clustering of the clinical samples showed two main groups with the major group characterized by high activities for s4 and low activities for s3 suggesting frequent WGD and consistent with previous observations^4,19^. The smaller group was predominantly composed of s1 mitotic errors and s3 HRD and may represent near diploid tumours. PDOs were well distributed across the two groups but there were three small subclusters which were underrepresented: those presenting a lack of s2 and s4, a lack of s2 and s3, and a lack of s3 together with high s4. PDOs derived from the same patient (PDO3 and PDO9, PDO5 and PDO6, and PDO7 and PDO8) were clustered together. Taken together, these data indicate that PDO represent the copy number mutational landscape observed in HGSOC patients.

### Effect of CNAs at the gene expression level

To understand how PDO absolute copy number alterations (CNAs) could alter the gene expression of corresponding genes, we first tested whether PDOs displayed known HGSOC-associated amplifications (Supplementary Fig. 8a) and which genes were highly amplified when averaged over all PDOs (Supplementary Fig. 8b), including the well characterized copy number drivers *MYC* and *CCNE1*. We performed RNAseq on the PDOs and compared their transcriptome to the TCGA primary tissue cohort and found highly similar cell-autonomous transcriptional profiles. As expected, we observed significant under-expression of genes relating to the tumour microenvironment (Fig. 2a) which is not represented in the organoid cultures. Principal component analysis on the scaled and centered DESeq2 counts showed that PDOs derived from the same patient PDO5 and PDO6 - the transcriptome of which is nearly identical - cluster together, but that PDO7 and PDO8, which are distinguished by a secondary *BRCA1* mutation following progression after PARP therapy, differ from each other (Fig. 2b). As PDOs consist of 100% tumour cells, we assessed the correlation between gene copy number changes and their expression using two metrics. The first metric shows whether, on average, PDOs with lower copy number values in genes have a lower gene expression, in order to capture nonlinear relationships between copy number and gene expression. We computed the average gene expression values for the three PDOs of lowest copy number and calculated the fraction of remaining PDOs with higher gene expression values than this average (Fig 2c). The second metric used was the R^2^ of the correlation between DESeq2 count values and absolute copy number in each gene across PDOs. For both metrics, higher values indicate stronger evidence for copy number driven gene expression (Fig. 2d). The most highly variable areas in the genome are located within chromosomes 8, 10, 11, 12, 17 and 1 (Supplementary Fig. 9a), where we found the most highly correlated genes. *MYC* showed the best good correlation between copy number and gene expression and was also the gene with the highest absolute copy number in our PDO cohort (Fig 2d), followed by *ZWINT* (Supplementary Fig. 9b).

**Fig. 2.**
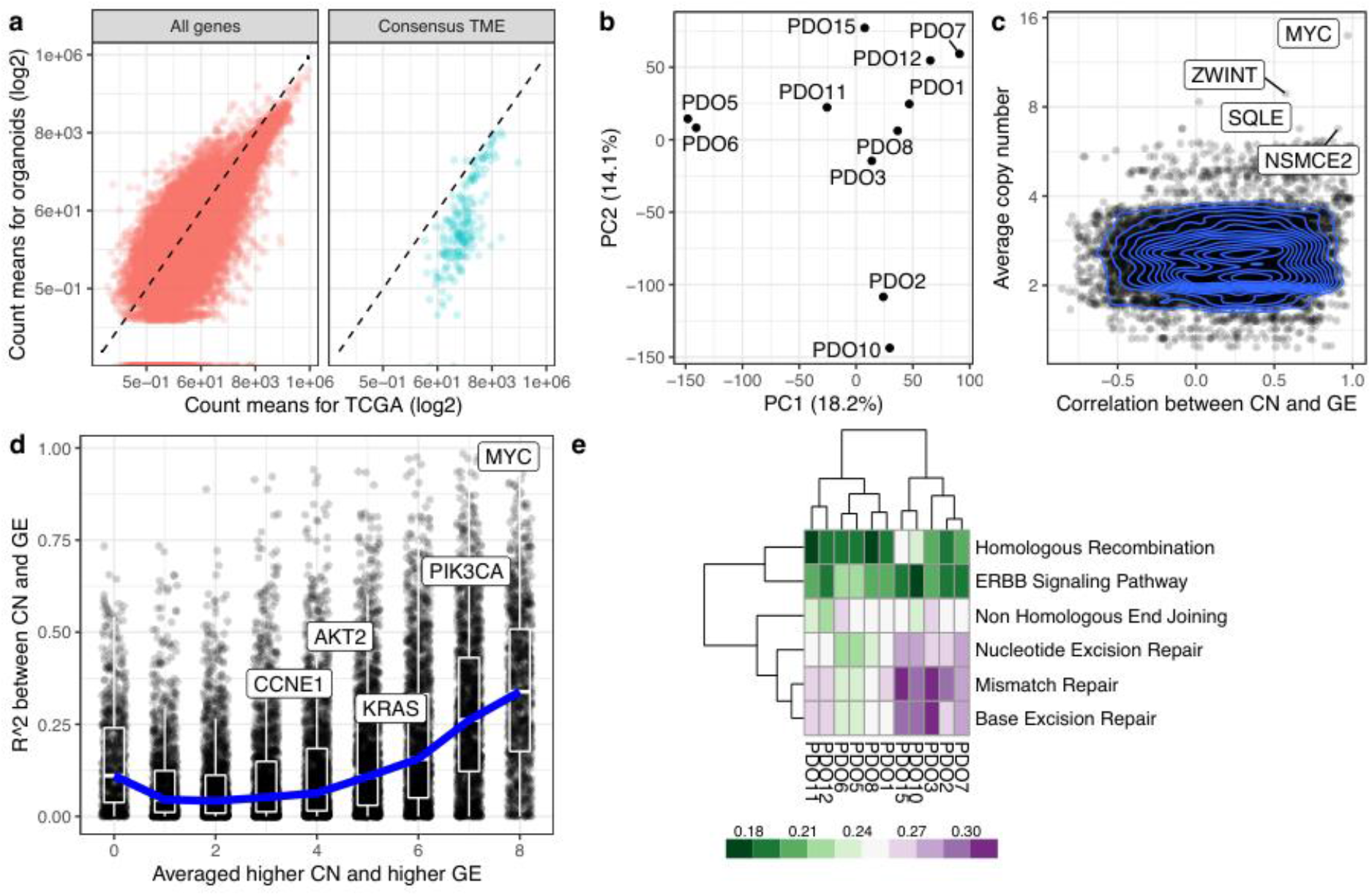
Transcriptomic analysis of HGSOC organoids **a** Scatterplots show correlation for the average counts, in transcripts per million (TPM) for each gene in the TCGA and the organoid cohorts. Consensus TME genes represent non-tumour genes expression in the tumour microenvironment^20^. The dashed line corresponds to the identity line. **b** Principal component analysis based on DESeq2 counts for 11 organoids. **c** Scatterplot and contour plot of the Pearson correlation coefficient for copy number and gene expression, and average absolute copy number for each gene. *MYC* and *ZWINT* are shown as highly correlated genes. **d** Scatterplot of two metrics for assessing the agreement between copy number and gene expression. For each gene, we computed the average expression of the three organoids with lowest copy number value. The metric is the fraction of remaining organoids which have higher gene expression value than this average, and takes values between 0/8 and 8/8, with higher values indicating greater agreement between copy number and gene expression between organoids. This is shown in the x axis. On the y axis we display the R^2^ value for the correlation between copy number state and gene expression. We have labelled genes of interest. The blue curve indicates the median R^2^ values in each group of the metric along the x-axis, and box-plots indicate the interquartile range. **e** DNA damage response pathway analysis from RNAseq on 11 PDOs. PDO10 and PDO15 show high enrichment scores for homologous recombination compared to other PDOs. Also mismatch and base excision repair pathways are high in these models.

As defects in DNA damage response pathways are clinically important for treatment, we tested for enrichment scores across the PDOs. PDO10 and PDO15 have a high enrichment score for homologous recombination deficiency (Fig. 2e), present nearly identical signature activities, and are the two PDOs with highest s7 activity (Fig. 1a).

### PDO drug screening

We compared drug sensitivity between PDOs and their parental uncultured patient-ascites. Using 13 anti-cancer compounds dispensed in an 8-point half-log dilution series, we found moderate to high correlation between the plasma drug concentration-time area under the curve (AUC) of PDO and their corresponding patient-derived ascites (Fig. 3a). We then tested all the PDOs using standard of care chemotherapy (oxaliplatin, paclitaxel, gemcitabine and doxorubicin) (Fig. 3b) as we observed no effect with the targeted therapies at the concentrations used in this study. Based on the median AUC we divided PDOs into two groups: sensitive (PDO1, PDO2, PDO3, PDO11, PDO12) and resistant (PDO5, PDO6, PDO7, PDO8, PDO10) (Supplementary Fig. 10) and performed differential gene expression and pathway analysis (Fig. 3c) to infer mechanisms of resistance. Sensitive PDOs showed increases in *MYC* targets and interferon alpha and gamma responses while resistant PDOs had an increase in hypoxia, *KRAS* signaling and epithelial-mesenchymal transition (EMT) pathways.

**Fig. 3.**
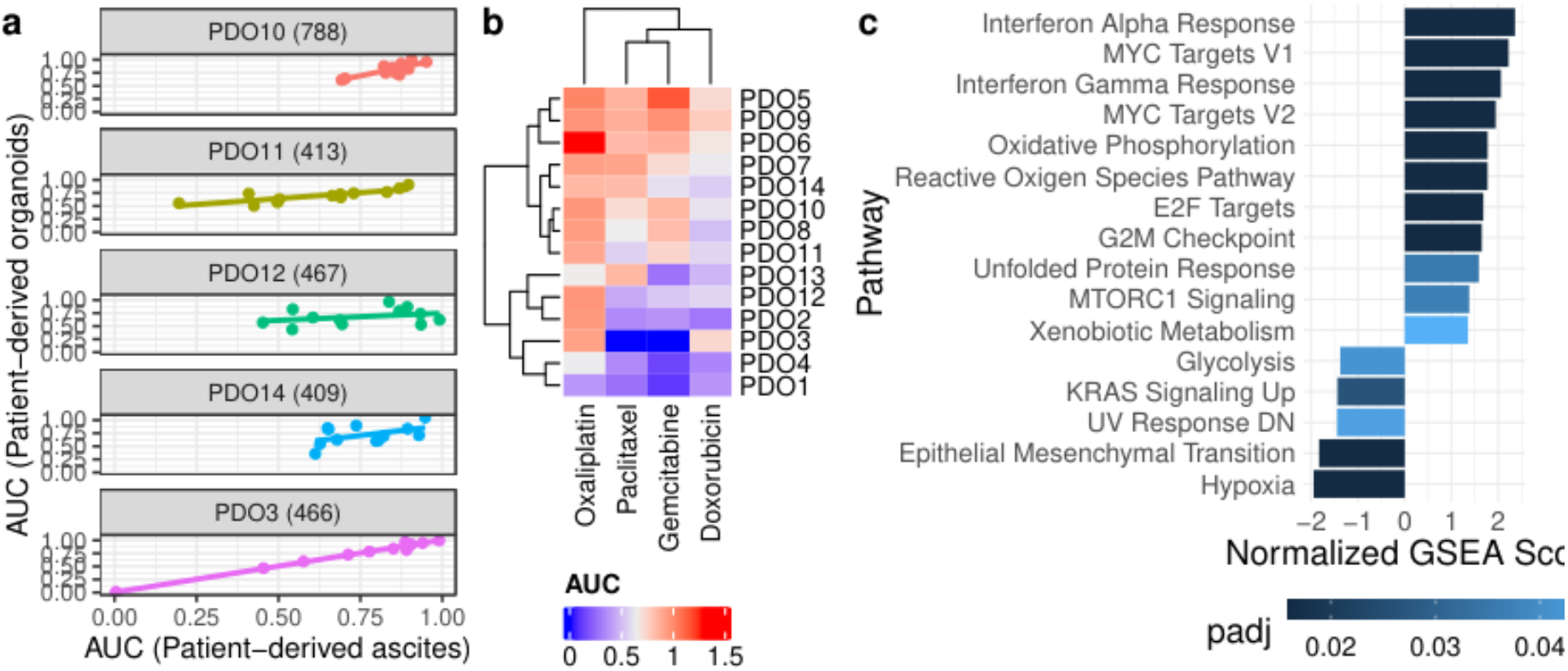
Organoids are clinically relevant models **a** Correlation of drug response between uncultured patient-ascites spheroids and their derived-PDOs using 13 compounds (PDO14: cor. 0.49, p-value 0.1; PDO11: cor. 0.82, p-value 0.001; PDO3: cor. 0.995, p-value 2.3e-11; PDO10: cor. 0.81, p-value 0.001; PDO12: cor.0.32, p-value 0.31). **b** Organoid drug responses to standard of care chemotherapies. The observed dose-response relationships were not always compatible with the Hill dose-response model assuming a sigmoidal decrease so that 5-parameter logistic model fits were preferred, explaining AUC estimates greater than 1. **c** Significant pathways based on adjusted p-value (padj) after performing Gene Set Enrichment Analysis (GSEA) with rank based on significance level, for the sensitive-to-resistant PDO comparison of intra-PDO heterogeneity.

### Organoid intratumoural heterogeneity

In order to assess genomic heterogeneity within PDOs, we performed single cell whole genome sequencing on three of the models: PDO2 (N = 76 cells), PDO3 (N = 145 cells) and PDO6 (N = 355 cells) (Fig. 4). Copy number changes at single-cell resolution revealed widespread clonal loss of heterozygosity (LOH) in large regions spanning up to entire chromosomes that were PDO specific (e.g. chromosome 13 in PDO6). Subclonal LOH, although less common, was also present in all three organoids. Amplification events were more common than losses; for example, chromosomes 2, 3, and 20 are clonally amplified in PDO2 and PDO3 whereas chromosomes 6 and 11 showed large, amplified regions shared between PDO3 and PDO6. All three PDOs present non-focal amplifications in chromosomes 1, 5, 12 and 20 as well as deletions in chromosome 13. This analysis also provided strong evidence for clonal amplification of candidate driver copy number aberrations: *CCNE1* in PDO2 and PDO3, an early chromothriptic event at *MYC* in PDO3 (Supplementary Fig. 11), and *AKT2* in PDO2 and PDO6. PDO6 showed early clonal loss of *RB1*.

**Fig. 4.**
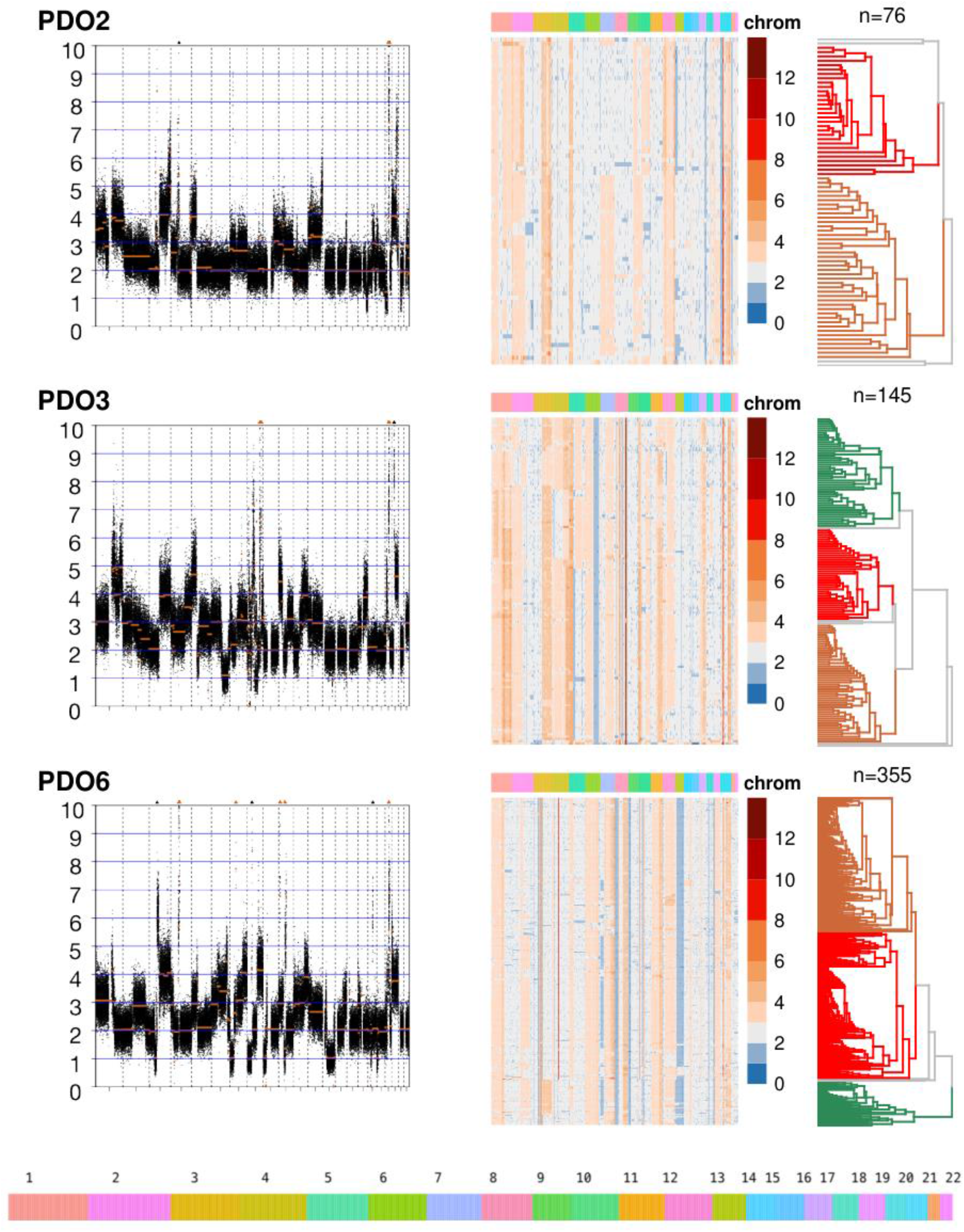
Genomic heterogeneity in 3 high grade serous carcinoma PDOs. The left column represents bulk absolute copy number profiles; the middle column shows single-cell DNA (scDNA) copy number where cells have been clustered using hierarchical clustering on Euclidean distance, and the right column shows absolute copy number density states for the scDNA in the indicated region (red arrow). Each row within the scDNA plots represents a cell across the different chromosomes in the x-axis and the copy number state (20Kb bins) is indicated in colours. Loss of heterozygosity and amplification events are common in all three patient-derived organoids. Examples of clonal populations within specific chromosomal regions of the organoids are shown, indicated by black boxes, for chromosome 7 in PDO2, for chromosome 5 in PDO3 and for chromosome 2 in PDO6 (same region in the bulk profiles is shown within a red box).

We also identified regions of clonal heterogeneity in all three PDOs (Fig. 4 and Supplementary Fig. 12). We quantified the heterogeneity observed in each PDO by comparing the observed copy number variance to the expected copy number variance (Methods), and found that, globally, PDO3 showed the highest subclonal heterogeneity, with 48% of the genome presenting subclonal heterogeneity, followed by PDO6 (29%) and PDO2 (26%) (Supplementary Fig. 13).

## Discussion

Our analysis of copy number features and mutational signatures shows that HGSOC PDOs recapitulate the broad mutational landscape of patient samples. The organoid models contained a mixture of signatures indicating the influence of multiple mutational processes. Although their copy number signatures are well spread across the range seen in patient samples, certain copy number combinations are underrepresented (high s4 and s7, high s3 and s5, and high s6). Critically, we show that PDO are also vital models to study heterogeneity at the single cell level and we found that, although all models tested showed genomic heterogeneity, the level of complexity varies. This suggests that different mutational processes may have different abilities to driving evolutionary change and PDO now provide tools for lineage tracing experiments to test this. Further analysis of clonal populations with PDO also has the potential to define the active mutational processes by sequential single-cell-cloning as recently described^21^. Lastly, these models also provide important insights into the genomic aetiology of HGSOC, including evidence for chromothripsis as an early initiation event in HGSC by targeting *MYC* and indicating that tandem duplication can occur in the absence of either *BRCA1* or *CDK12* mutation.

The development of high quality pre-clinical tumour models is of high importance for therapeutic discovery in HGSOC. Existing cell-based and PDX models have not been characterized in detail and their relationship to the diversity of CIN seen in patient tissue samples is unknown. Derivation of continuous cell lines has proven difficult for HGSOC, and although new cell lines are being developed^22–24^ success rates are comparatively low and the number of available models has not significantly increased over the past 10 years. With the wider use of organoid culture, ovarian cancer models have been developed both as short and long-term cultures^10–13^ but with variable information about success rates and survival in culture. We demonstrated that short-term HGSOC organoid derivation from human ascites samples and PDX tissues can be achieved with good efficiency. However, as indicated by our time to event analyses, further improvements in media and culture conditions are needed to improve success rates, particularly from solid tissue samples.

Although SCNAs have been shown to affect gene expression levels for the most abundantly expressed human genes indicating global gene dosage sensitivity^25^, it has also been described that this correlation does not always translate proportionally due to transcriptional adaptive mechanisms^26^. In our study we compared PDO gene expression to TCGA patient samples and corroborated that gene transcript levels are highly correlated, providing ideal models to study tumour cell intrinsic associations. We have previously found that the correlation between SCNA and gene expression is higher for cancer driver genes that are frequently amplified and identified co-dependencies between amplification of *MYC* and genes from the *PI3K* pathway which have therapeutic potential^27^. We corroborated, using novel ways of correlating absolute SCNA with transcriptomics, that in our organoid models, correlation was highest for *MYC*, *PIK3CA* and *AKT2* reinforcing their putative role as potential targetable cancer drivers.

Genetic alterations in HGSOC are extraordinarily diverse therefore the development of a truly personalised treatment requires genomically annotated individual patient avatars for therapeutics. In this study, we showed the potential of HGSOC PDOs as a new preclinical cancer model representing individual patients. Consistent with studies in ovarian cancer and other tissue types^9,28–30^ our results confirm the feasibility of using PDOs for testing drug sensitivity in HGSOC. Future studies should account for doubling-time confounding errors using different metrics such as Growth Rate (GR) metrics^31^.

This study has shown that HGSOC PDOs faithfully represent the high variability in copy number genotypes observed in HGSOC patients and together with their associated clinical, phenotypic and genomic characterizations will provide an important resource for pre-clinical and translational studies investigating genomic biomarkers for treatment stratification and to further our understanding of tumour heterogeneity and clonality.

## Methods

### Ethical approval and clinical data collection

Clinical data and tissue samples for the patients were collected on the prospective cohort study Cambridge Translational Cancer Research Ovarian Study 04 (CTCR-OV04) and was approved by the Institutional Ethics Committee (REC08/H0306/61). Patients provided written, informed consent for participation in this study and for the use of their donated tissue for the laboratory studies carried out in this work. Clinical data for all the patients is provided in Supplementary Information.

### Sample collection and processing

Samples were obtained from surgical resection, therapeutic drainage or surgical washings. Solid tumours were assessed by a pathologist and only tumour samples with ≥ 50% cellularity were attempted to grow. A small portion of each sample was kept at −80°C until used for genomic profiling.

### Organoid derivation

Tumour samples were washed in PBS, minced into 2mm pieces using scalpels and incubated with gentamicin (50μg/ml), Bovine Serum Albumin Fraction V (1.5%), insulin (5μg/mL), collagenase A (1 mg/mL) and hyaluronidase (100U/ml) for 1–2h at 37°C. Following incubation, the mixture was filtered and the cell suspension was spun down and washed with PBS. Ascites fluid was centrifuged at 450g for 5 min. Cells were then washed with PBS and centrifuged at 400g for 5 min.

The isolated cells were resuspended in 7.5 mg/ml basement membrane matrix (Cultrex BME RGF type 2 (BME-2), Amsbio) supplemented with complete media and plated as 20 μl droplets in a 6-well plate. After allowing the BME-2 to polymerize, complete media was added and the cells left at 37 °C. We used published culture conditions for normal fallopian tube growth14 as follows: AdDMEM/F12 medium supplemented with HEPES (1×,Invitrogen), Glutamax (1×, Invitrogen), penicillin/streptomycin (1×, Invitrogen), B27 (1×, Invitrogen), N2 (1×, Invitrogen), Wnt3a-conditioned medium (25% v/v), RSPO1-conditioned medium (25% v/v), recombinant Noggin protein (100 ng/ml, Peprotech), epidermal growth factor (EGF, 10 ng/ml, Peprotech), fibroblast growth factor 10 (FGF10, 100 ng/ml, Peprotech), nicotinamide (1 mM, Sigma), SB431542 (0.5 μM, Cambridge Biosciences), and Y27632 (9 μM, Abmole).

### Organoid culture

Organoid culture medium was refreshed every 2 days. To passage the organoids, the domes were scraped and collected in a falcon tube, TrypLE (Invitrogen) was added and incubated at 37 °C for approximately 10 min. The suspension was centrifuged at 800g for 2 min and the cell pellet was resuspended in 7.5 mg/ml BME-2 supplemented with complete media and plated as 20 μl droplets in a 6-well plate. After allowing the BME-2 to polymerize, complete media was added, and cells incubated at 37 °C. The commonest cause of culture failure was growth arrest or fibroblast overgrowth. We considered an organoid line to be continuously established when it had been serially passaged >5 times followed by cryopreservation and successful re-culture. By these criteria, 15/18 PDO lines were continuous.

### Immunohistochemistry

Haematoxylin and Eosin (H&E) slides were stained according to the Harris H&E staining protocol and using a Leica ST5020 multi-stainer instrument. Paraffin embedded sections of 3 μm were stained using Leica Bond Max fully automated IHC system. Briefly, slides were retrieved using sodium citrate for 30 minutes and p53 antibody (D07, 1:1000, Dako) was applied for 30 minutes. Bond™ Polymer Refine Detection System (Leica Microsystems) was used to visualise the brown precipitate from the chromogenic substrate, 3,3’-Diaminobenzidine tetrahydrochloride (DAB).

### Nucleic acid isolation

DNA and RNA were extracted at the same time from the same cells. Extraction was performed using the DNeasy Blood & Tissue Kit (QIAGEN) according to manufacturer instructions.

### Bulk shallow whole-genome sequencing

Whole genome libraries were prepared using the TruSeq Nano Kit according to manufacturer instructions. Each library was quantified using the KAPA Library Quantification kit (kappa Biosystems) and 10nM of each library was combined in a pool of 21 samples and sequenced on the Illumina HiSeq 4000 machine using single-end 150-bp reads. Reads were aligned against the human genome assembly GRCh37 using the BWA-MEM algorithm (v0.7.12). Duplicates were marked using the Picard Tool (v1.47) and copy number was assessed using the Bioconductor package QDNAseq (v1.6.1)^33^.

### Absolute copy number signature analysis

Copy number signatures for the organoid cultures were calculated as previously described^6^.

### TCGA, Britroc and PCWAG

Signature activities of organoids were compared to those previously described in three HGSOC cohorts: TCGA and Britroc^6^ (sWGS-based signatures) and PCAWG^4^ (WGS-based signatures). Copy number signature activities were transformed using the centered log-ratio transformation with an imputation value of 10^−2^ to consider that they are compositional data which sample-wise add up to one. Organoid and primary tissue samples were clustered using hierarchical clustering with complete linkage on this transformed space. We performed additional analyses to confirm that our conclusions – namely, that the signature activities of organoids are representative of the activities of primary tissue, and in determining which activities are underrepresented in the organoids – were robust to the imputation value. Using imputation values between 0.001 and 0.1 we show that the dendrogram in Figure 1 is similar to the dendrograms generated using both higher and lower imputation values, and that the underrepresented clades are robust to changes in the imputation values. A more detailed report of the differences in dendrograms as we vary the imputation values can be found in the Github repository.

### Single cell shallow whole genome sequencing

Organoids were dissociated into single cells using TrypLE, washed twice with PBS and counted. Single cell solution was filtered using a 70μm Flowmi^®^ filter to remove any duplets or triplets. With the aim of getting around 300 cells for library preparation, 4000 single cells were loaded on the chip. Single cell 10x CNV libraries were prepared according to the manufacturer’s protocol (10X Genomics) and multiplexed in equal molarity to achieve 2.4 million reads per cell. Single cell 10X CNV constructed libraries were sequenced on Illumina Novaseq6000 S4 platform using PE-150 mode.

### Metric for copy number subclonal heterogeneity in single cell

The metric for copy number subclonal heterogeneity is defined as follows. Independently, for each of the three organoids, we fitted a linear model of the standard deviation of the absolute copy number across organoids predicted by its mean, using bins of 500kb. Copy number data were handled using the R package GenomicRanges^34^. The marked positive correlation indicated that the data were heteroscedastic. For each bin we computed its expected variance from the model, *E*(*σ*^2^), and compared it to the observed variance *S*^2^ with a Chi-Squared test with alternative hypothesis *E*(*σ*^2^) < *S*^2^. A statistically significant result indicates that we see a greater variance than expected in the copy number values of this bin, and that therefore there is subclonal heterogeneity.

### Clade analysis of single cell copy number data

Single cell clades for each organoid were identified by performing hierarchical clustering using complete linkage on euclidean distance of copy number values on 500 kb-binned genomes. Only clades with more than 3 cells were kept in the analysis. PDO2 had four major clades, two of which encompassing most cells (clade A: 42 cells, clade B: 30 cells), PDO3 had 7 major clades, three of which with more than two cells (clade A: 40 cells, clade B: 52 cells, clade C: 48 cells). PDO6 had six clades, three of which contained more than one cell (clade A: 158 cells, clade B: 145 cells, clade C: 49 cells). The copy number profile comparison of the two clades of PDO2, and of the two pairwise comparisons of PDO3 and PDO4, were carried out using the 20 kb-binned copy number profile. Bins of distinct copy number between cells in different clades were detected using a Holm-Bonferroni-adjusted t-test on the absolute copy number value.

### Tagged-amplicon sequencing

Coding sequences of *TP53, PTEN, NF1, BRCA1, BRCA2, MLH1, MSH2, MSH6, PMS2, RAD51C, RAD51B, RAD51D*, and hot spots for *EGFR*, *KRAS*, *BRAF*, *PIK3CA* were sequenced using tagged amplicon sequencing on the Fluidigm Access Array 48.48 platform as previously described30. Libraries were sequenced on the MiSeq platform using paired-end 125bp reads. Variant calling from sequencing data was performed using an in-house analysis pipeline and IGV software^36^.

### RNAseq

RNA quality control was performed using Tapestation according to manufacturer’s and samples were processed using Illumina’s TruSeq stranded mRNA kit with 12 PCR cycles according to manufacturer’s instructions. Quality control of libraries was performed using Tapestation and Clariostar before normalising and pooling. Samples were sequenced using two lanes of SE50 on a HiSeq 4000 instrument. The analysis was performed using an in-house DESeq237 pipeline.

TCGA gene expression values were downloaded as HTSeq count files of Genome Build GRCh38 for 240 ovarian samples of either progressive disease, or complete remission or response. The counts were normalised using DESeq2. The subset of genes relating to the tumour microenvironment were taken from the Consensus^TME^ list (https://github.com/cansysbio/ConsensusTME).

### Pathway enrichment analysis

Using our transcriptomic data, we computed enrichment scores for KEGG pathways of interest using ssGSEA, implemented in the R package gsva^38^, and using gene sets from the package GSVAdata^38^. To determine which pathways were overrepresented in the differential expression analysis between sensitive and resistant samples we used the R package fgsea^39^ and selected the top ten pathways according to their adjusted p-value (Benjamini-Hochberg correction), using the Hallmark gene sets from MSigDBv5p2.

### Drug sensitivity

An 8-point half-log dilution series of each compound was dispensed into 384 well plates using an Echo^®^ 550 acoustic liquid handler instrument (Labcyte) and kept at −20°C until used. Prior to use plates were spin down and 50 μl of organoid suspension is added per well using a Multidrop™ Combi Reagent Dispenser (Thermo-Fisher). Following 5 days of drug incubation cell viability was assayed using 30μl of CellTiter-Glo^®^ (Promega). Screens were performed in technical triplicate.

Drug response measures were standardised by dividing the original values by the median drug response observed in the control group of each drug and sample and then modelled as a function of the dose (on the log scale) by means of a 4th degree polynomial robust regression, fitted by means of the function lmrob of the R package robustbase^40^. Drug response measures that obtained robust weights smaller than 0.4 (out of a range which spreads from 0 for outliers to 1 for non-outliers) were considered as outliers. After excluding outliers, we modelled the standardised drug response measures as a function of the dose (on the log scale) by means of five-parameter log-logistic model (drm function of the drc R package^41^ with fct argument set to LL2.5). Area under the curve estimates were finally obtained by integrating the expected standardised drug response given the dose on the dose range of interest (on the log scale). Note that the use of M-splines instead of a log-logistic model led to similar AUC estimates.

Compounds used in this study included standard of care chemotherapeutics paclitaxel (Sigma), oxaliplatin (Selleck), doxorubicin (Selleck), and gemcitabine (Selleck). Maximum drug concentration in the assay was 30μM apart from paclitaxel (0.3μM) and oxaliplatin (300μM).

### *In vivo* growth

Animal procedures were conducted in accordance with the local AWERB, NACWO and UK Home Office regulations (Animals Scientific Procedures Act 1986). 1.5×10^5^ organoids were resuspended in 150 μl of PBS and injected intraperitoneally into NOD-scid IL2Rγ(null) (NSG) mice. Tumour growth was monitored by palpation and weighing the mice weekly.

## Data availability

RNASeq data are available at the Gene Expression Omnibus (GEO) under accession number GSE208216, and sWGS and scDNA data are available at the EGA European Genome-Phenome Archive (EGA) under accession number to be confirmed.

## Code availability

All the analysis code is at https://github.com/lm687/Organoids_Compositional_Analysis.

## Author Contributions

M.V., L.M.G. and J.D.B. wrote the manuscript. M.V. performed all the experiments. M.V., L.M.G, G.M., D-L.C. S.B., B.H. and C.M.S. performed data analysis. M.V., and D.S. derived and maintained the organoids. J.H. maintained the organoids. A.M.P. performed single cell sequencing. F.C-M. helped curate clinical data. F.M., G.M. and J.D.B. supervised the work.

## Acknowledgements

We thank all patients who participated in and donated tissue samples to this study. The Addenbrooke’s Human Research Tissue Bank is supported by the NIHR Cambridge Biomedical Research Centre. L.M.G. was supported by the Wellcome Trust PhD programme in Mathematical Genomics and Medicine (grant number RG92770). We also thank Karen Hosking, Mercedes Jimenez-Linan, and the OV04 study team for their help with clinical tissue samples. We would like to thank the Cancer Research UK Cambridge Institute Genomics, IT & Scientific Computing, Biological Resource Unit, Compliance & Biobanking, Research Instrumentation and Cell Services, and Bioinformatics core facilities for their support with various aspects of this study. The results shown here are in part based upon data generated by the TCGA Research Network: https://www.cancer.gov/tcga. We acknowledge funding and support from Cancer Research UK and the Cancer Research UK Cambridge Centre (A22905, A29580, A25117). This research was also supported by the NIHR Cambridge Biomedical Research Centre (BRC-1215-20014). The views expressed are those of the authors and not necessarily those of the NIHR or the Department of Health and Social Care. We thank Susana Ros, Thomas Bradley and Hayley Frances for critically reading the manuscript. We would like to thank the Clevers Laboratory (University of Utrecht) for hosting Maria Vias for a CRUK Travel Award.

## Supplementary Figures

**Supplementary Fig. 1.**
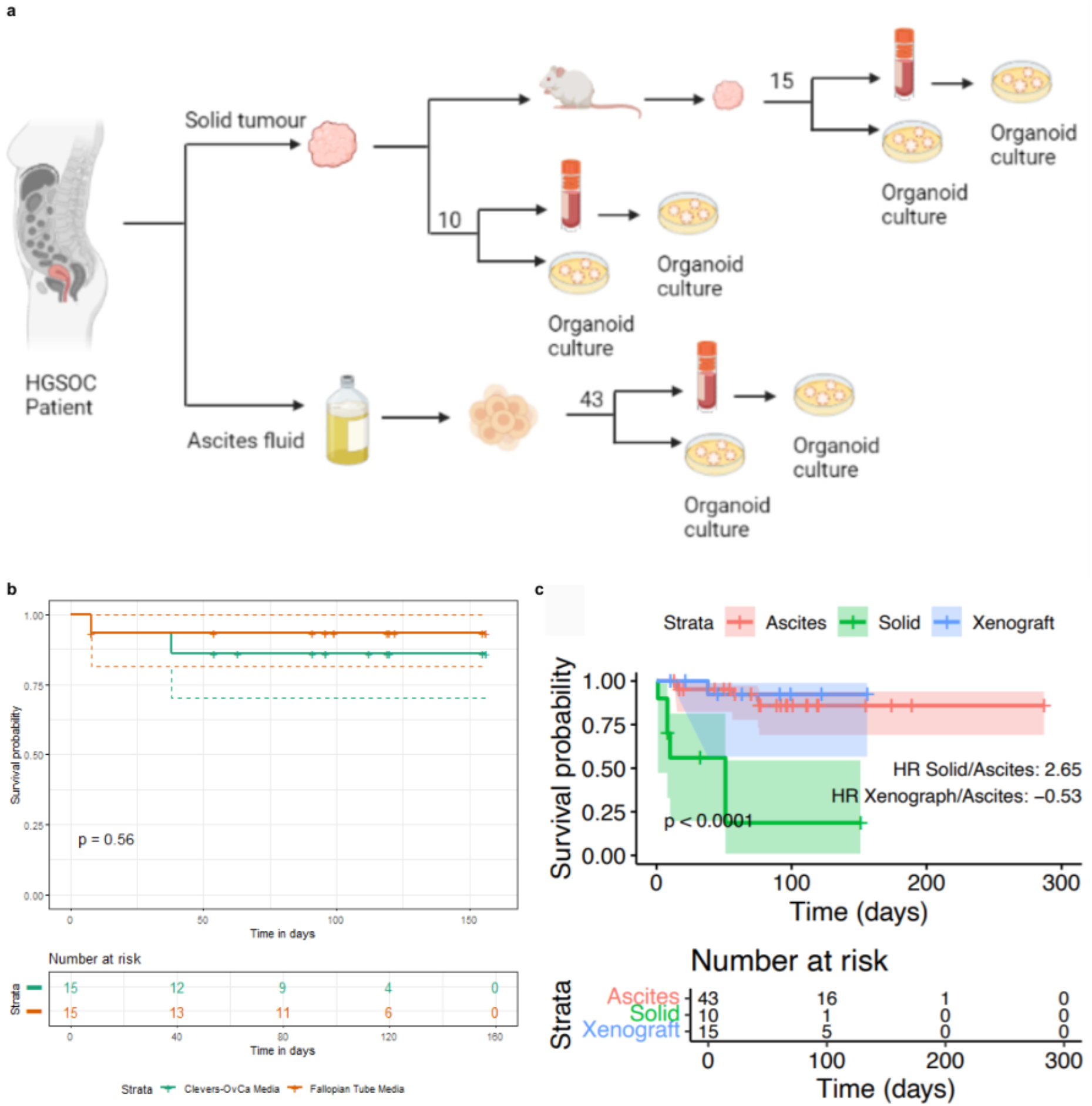
Sample collection workflow and survival analysis a Schematic of the sample collection workflow used in this study. b Survival analysis based on the type of media used to grow organoids. Two media were tested: formulation used in the Clevers lab for lab for growing ovarian cancer tissues and formulation used to grow fallopian tube tissue in the Meyers’s lab. P value was derived from the log rank test. **c** Kaplan-Meier survival curves showing association between type of tissue sample used for organoid derivation and survival of cultures. Survival probability is displayed as a function of time in days. Shading indicates the 95% confidence interval for each group. Hazard ratio and p-value were obtained from a log-rank test. Crosses correspond to censored observations.

**Supplementary Fig. 2.**
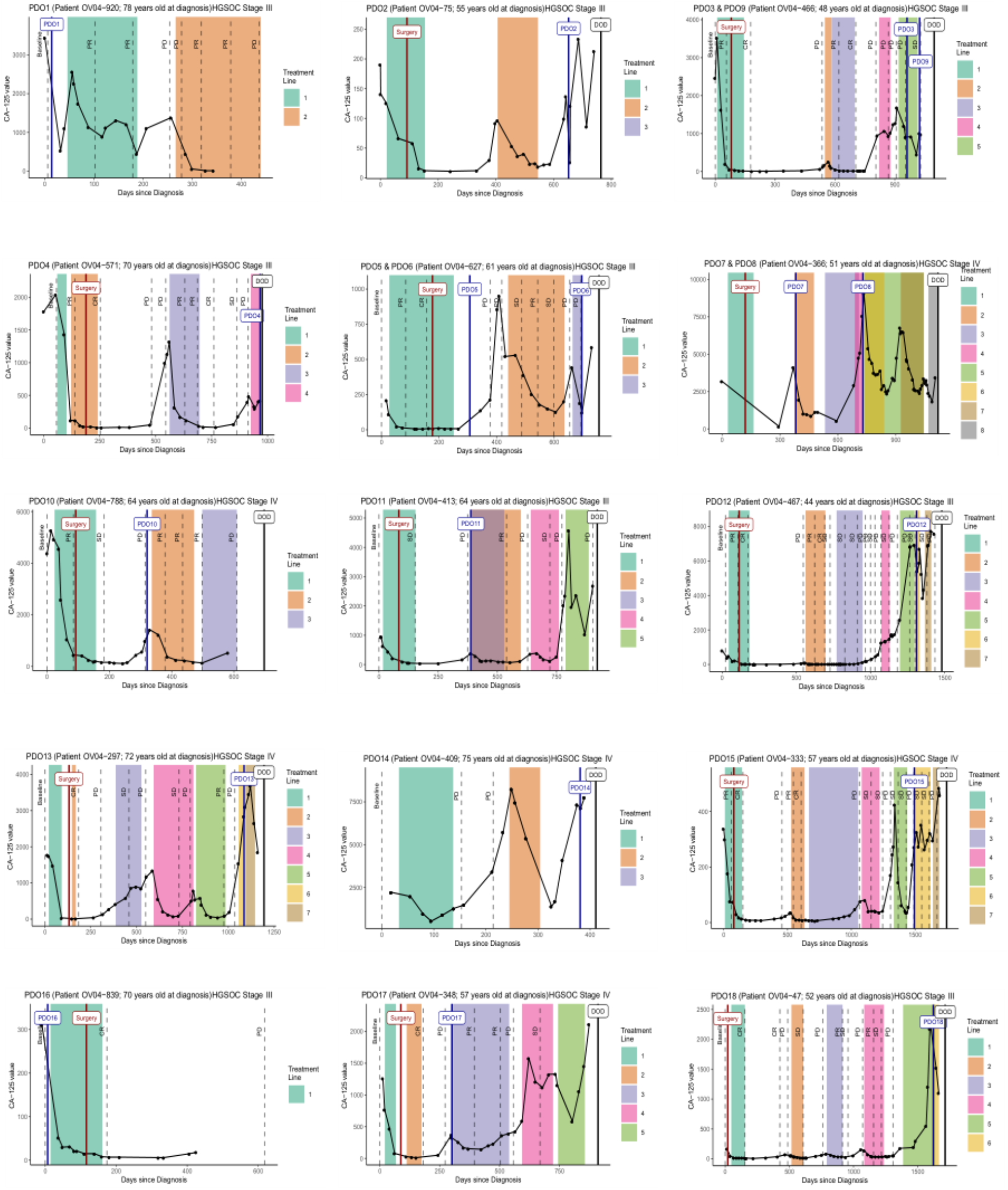
Clinical data Patient timeline summarising clinical data: CA125 levels, chemotherapy regimens and computerized tomography (CT) scans. CT scan outcomes are represented as follows: Baseline as first scan before treatment, PR as partial response, CR as complete response, SD as stable disease and PD as progressive disease. Treatment lines are represented in different colours (information on the specific treatment regimens is available in Supplementary Table 1) and patient date of death is shown as DOD. Date of ascites collection for organoid derivation is shown with the name of the organoid.

**Supplementary Fig. 3.**
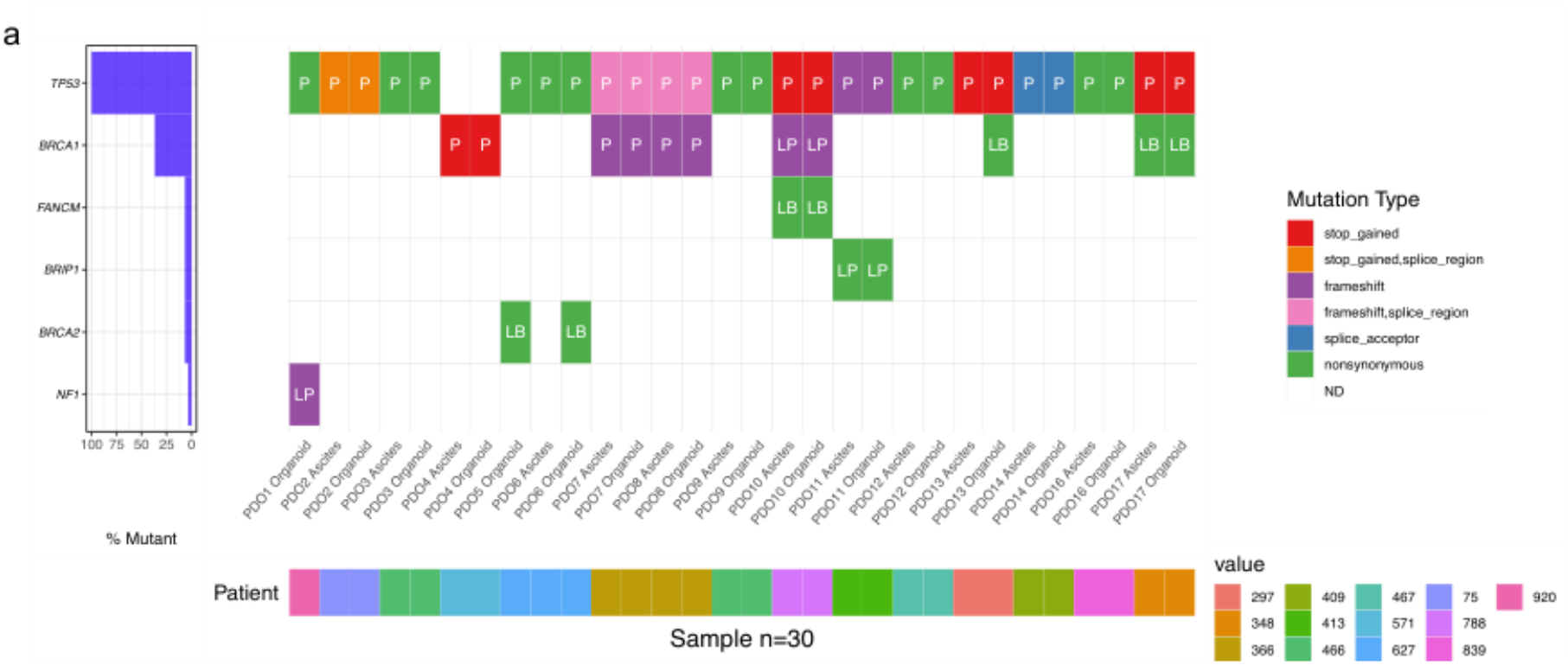
| Mutation analysis of PDO and patient samples. a Oncoplot showing mutation status on a specific gene panel. P represents pathogenic mutation, LP likely-pathogenic, and LB likely-benign.

**Supplementary Fig. 4.**
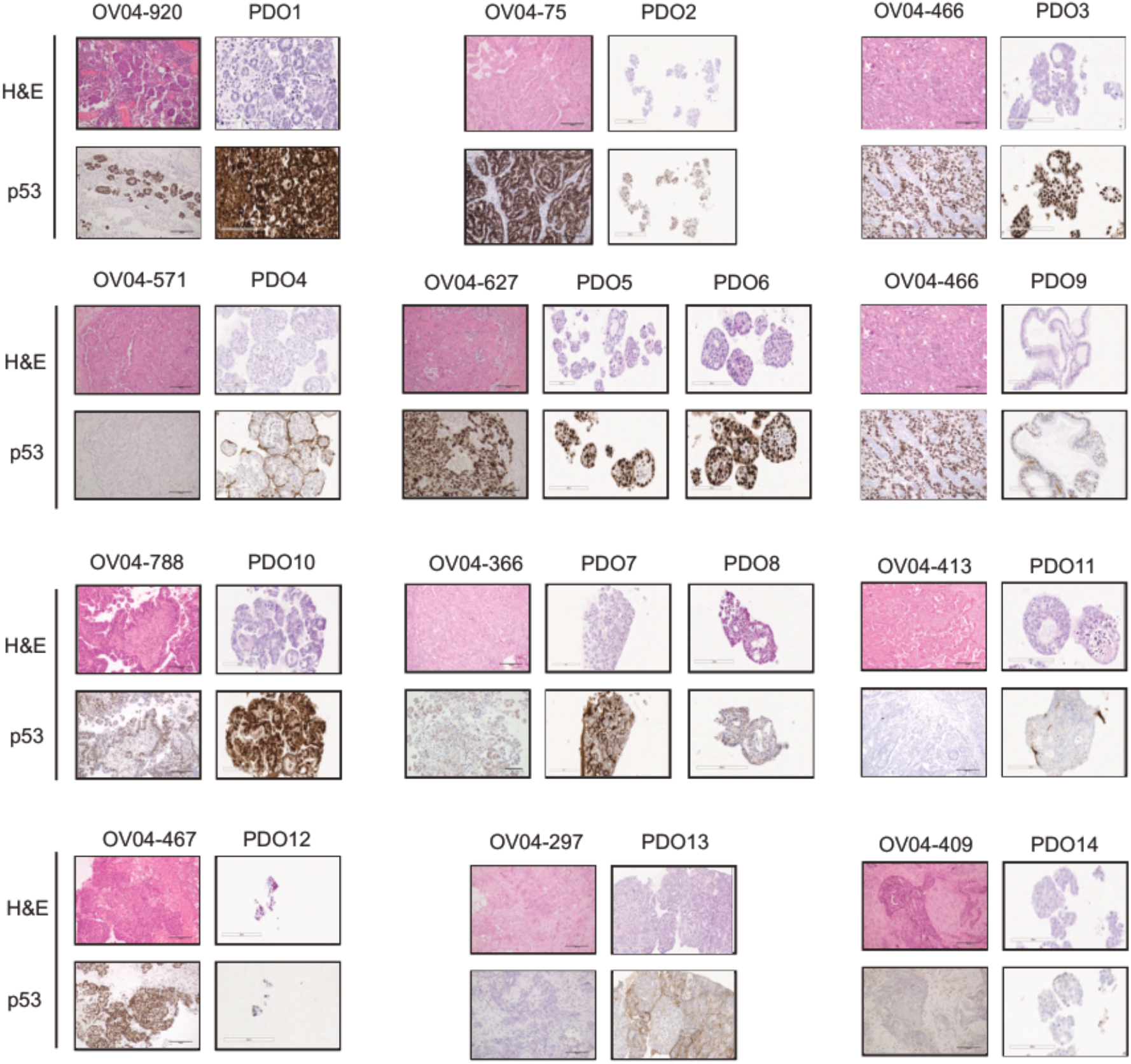
Tissue and PDOs morphological structures and p53 status. Tissues and PDOs sections were both Hematoxylin and Eosin and p53 stained. High grade serous ovarian carcinoma tissues contain specific structures as indicated by arrows that PDOs mimicked. The mutant p53 prevalence observed in HGSOC patients was reflected in the organoids with only one model displaying wild-type p53 expression patterns, some with loss expression and most of them with intense nuclear staining. PDO1 also shows similar glandular and micropapillary growth patterns. Microscope lens with 20× magnification was used for both tissues and organoids. Scale bars= 200 μm.

**Supplementary Fig. 5.**
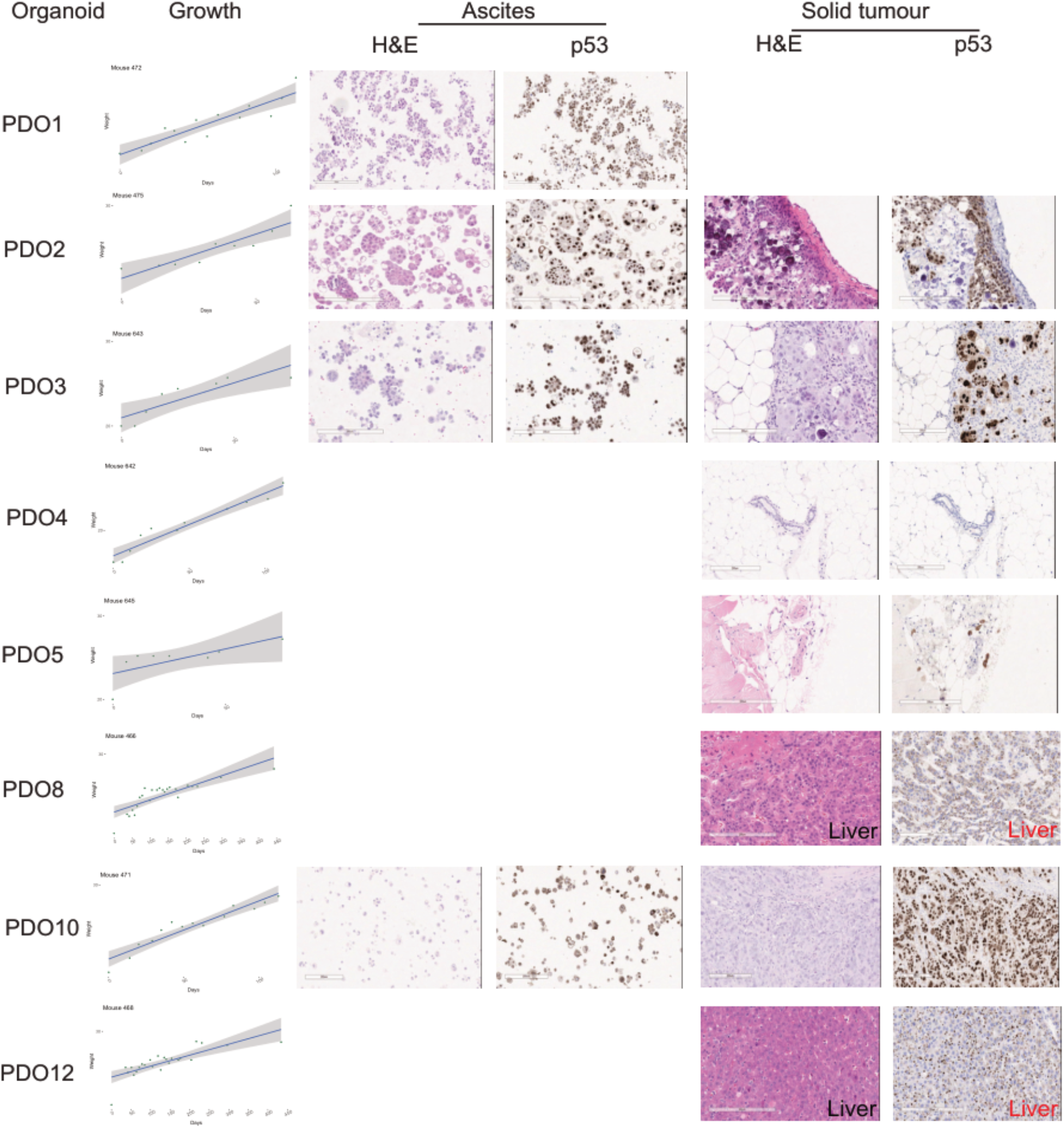
Orthotopic implantation of PDO. PDOs were implanted peritoneally into immunodeficient mice and disease progression was monitored by weighing the mice. Some mice developed ascites while others had disease in the liver or peritoneum. H&E and p53 immunostained sections are shown for each tissue collected. Scale bars = 200μm.

**Supplementary Fig. 6.**
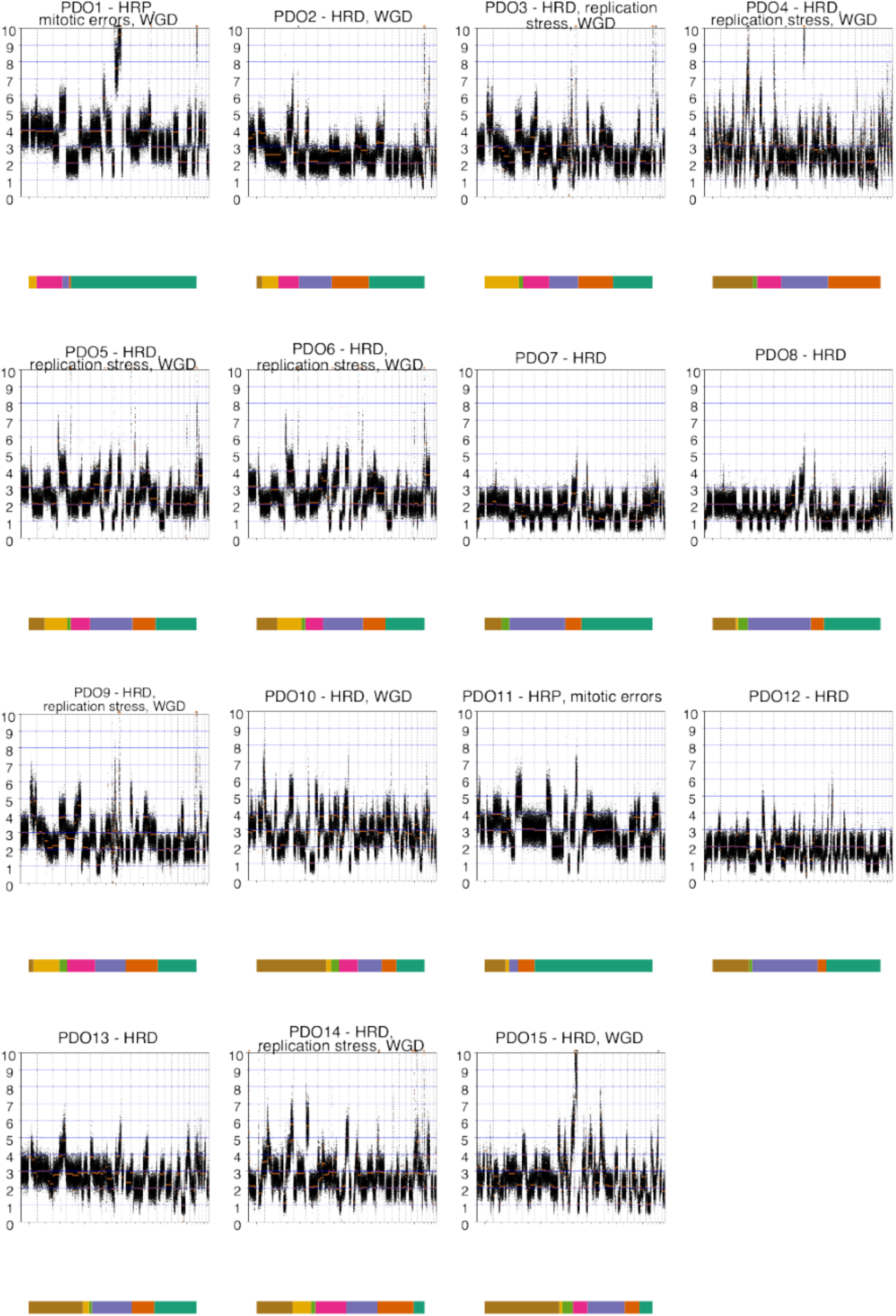
PDOs genome-wide absolute copy number alteration analysis. Absolute copy number profiles at 30Kb bin size are presented for all the PDOs. (PDO16, 17 and 18 were not continuous models.)

**Supplementary Fig. 7.**
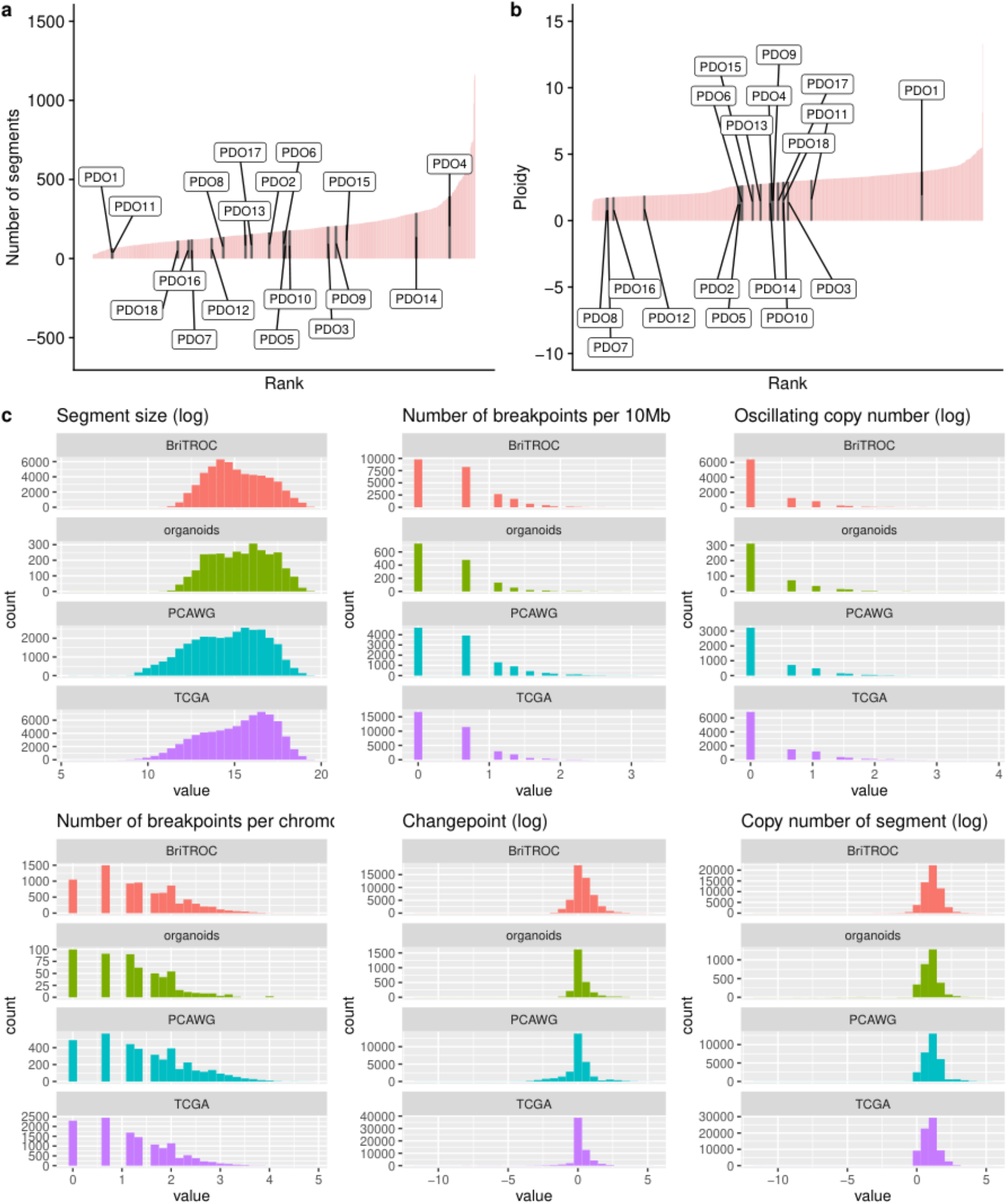
Comparison of genomic features between PDO and HGSOC cases in public data sets **a** and **b** Ordered barplots show the distribution of the number of copy number events and ploidy for organoids (labelled black bars) and 692 HGSOC tumours from the publicly available datasets of TCGA, PCAWG and BriTROC-1 (pink bars) c Tumour datasets used for comparison to PDOs (green bars) included TCGA (purple), BriTROC (orange) and PCAWG (blue). All genomic features are presented in log scale. Welch Two Sample t-test p-value on log-transformed data: number of breakpoints per 10MB (p = 0.61), segment size (p = 0.45), oscillating copy number (p = 0.72), number of breakpoints per chromosome arm (p = 0.60), number of changepoints (p = 0.85), and the copy number of the segments (p = 0.73).

**Supplementary Fig. 8.**
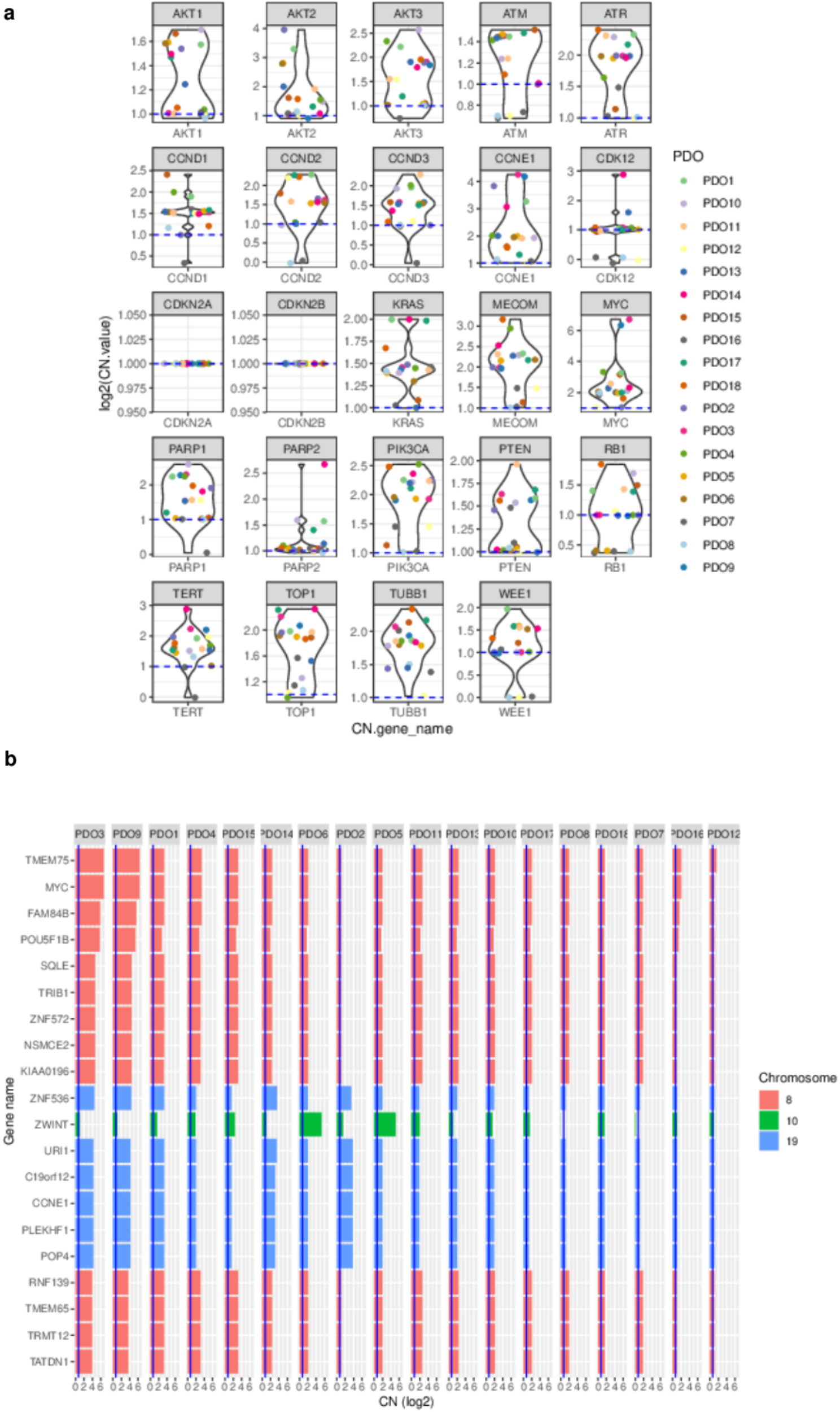
Absolute gene copy number in PDOs. a Absolute gene copy number for a set of important high grade serous ovarian cancer genes. b Absolute gene copy number for the most amplified genes when averaged across all the organoids.

**Supplementary Fig. 9.**
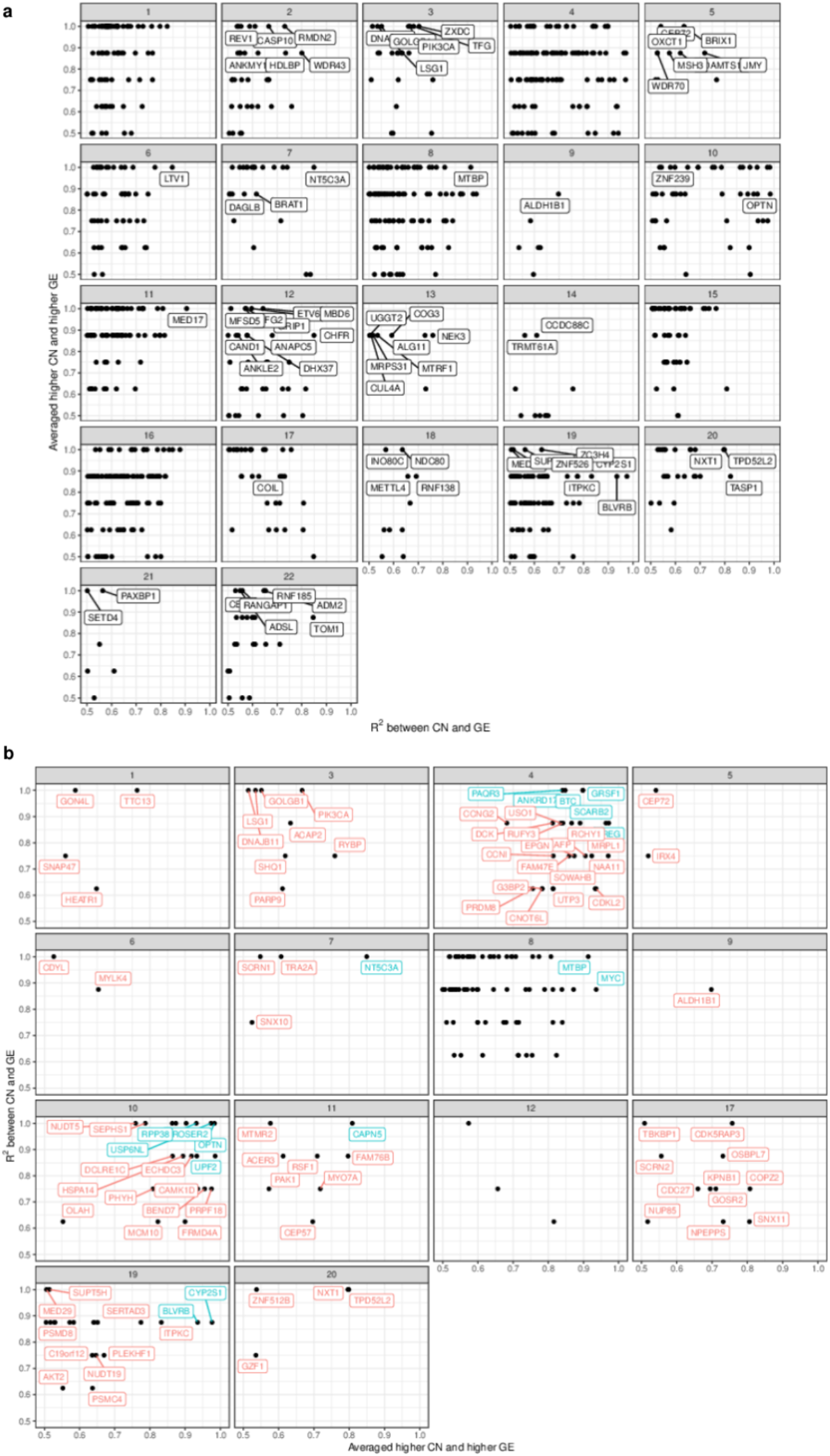
Whole genome correlation between absolute gene copy number and expression. a Scatterplot assessing the agreement between copy number and gene expression in genes of high copy number variability across different chromosomes. For each gene, we computed the average expression of the three organoids with lowest copy number value where the metric is the fraction of remaining organoids which have higher gene expression value than this average. A second metric we computed is the R^2 value. In both cases higher values indicate greater agreement between copy number and gene expression between organoids across different chromosomes. Chromosomes 14, 15, 16, 18 and 21 had no variable regions. b Genes with the highest values for both metrics have been labelled in red across different chromosomes and the top 20 are labelled in blue. Chromosomes 2 and 13 contained highly variable regions with low correlation.

**Supplementary Fig. 10.**
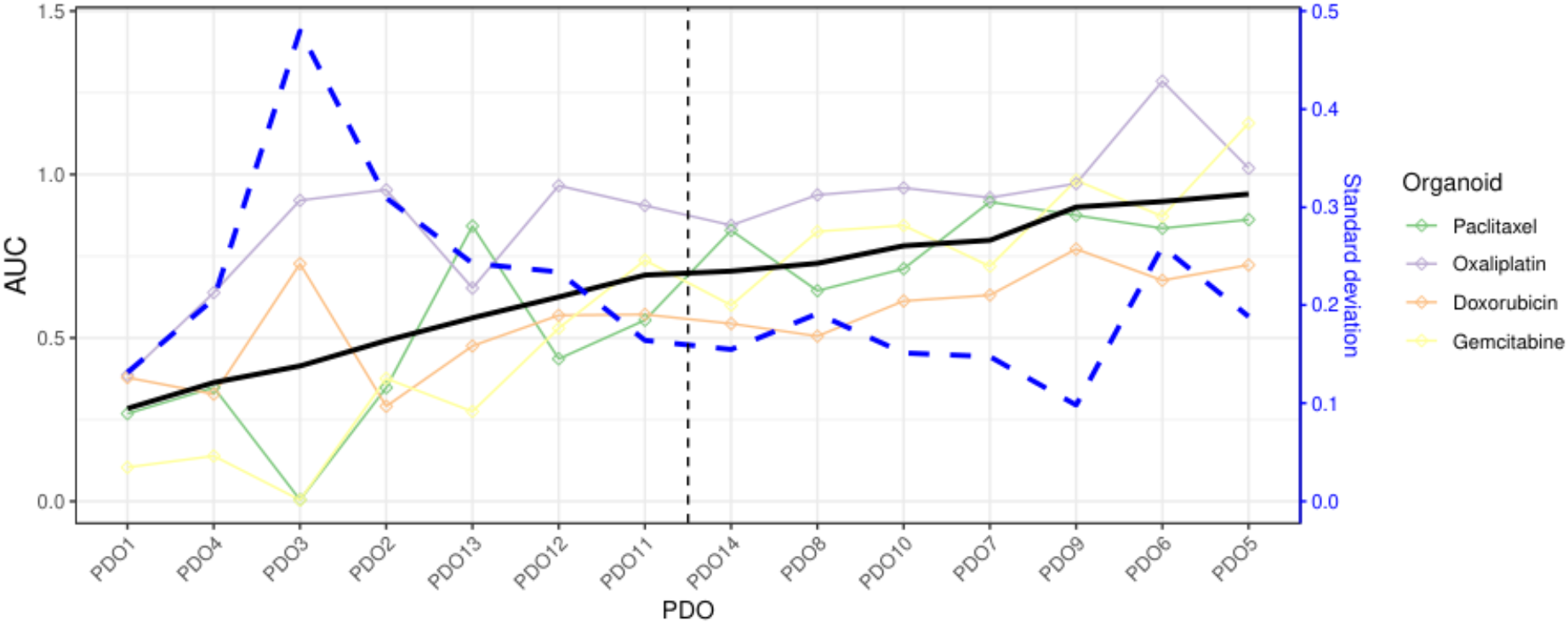
PDOs can be classified in two groups according to their drug sensitivity. Plot shows AUC (y-axis) of each drug (lines and points shape) for each sample (x-axis) ordered in increasing mean AUC values. Black line reports the mean AUC and the vertical dotted line suggests a split by mean AUC. The blue dashed line indicates the standard deviation of AUC. We grouped the samples into resistant and sensitive based on the mean AUC for our differential gene expression analysis.

**Supplementary Fig. 11.**
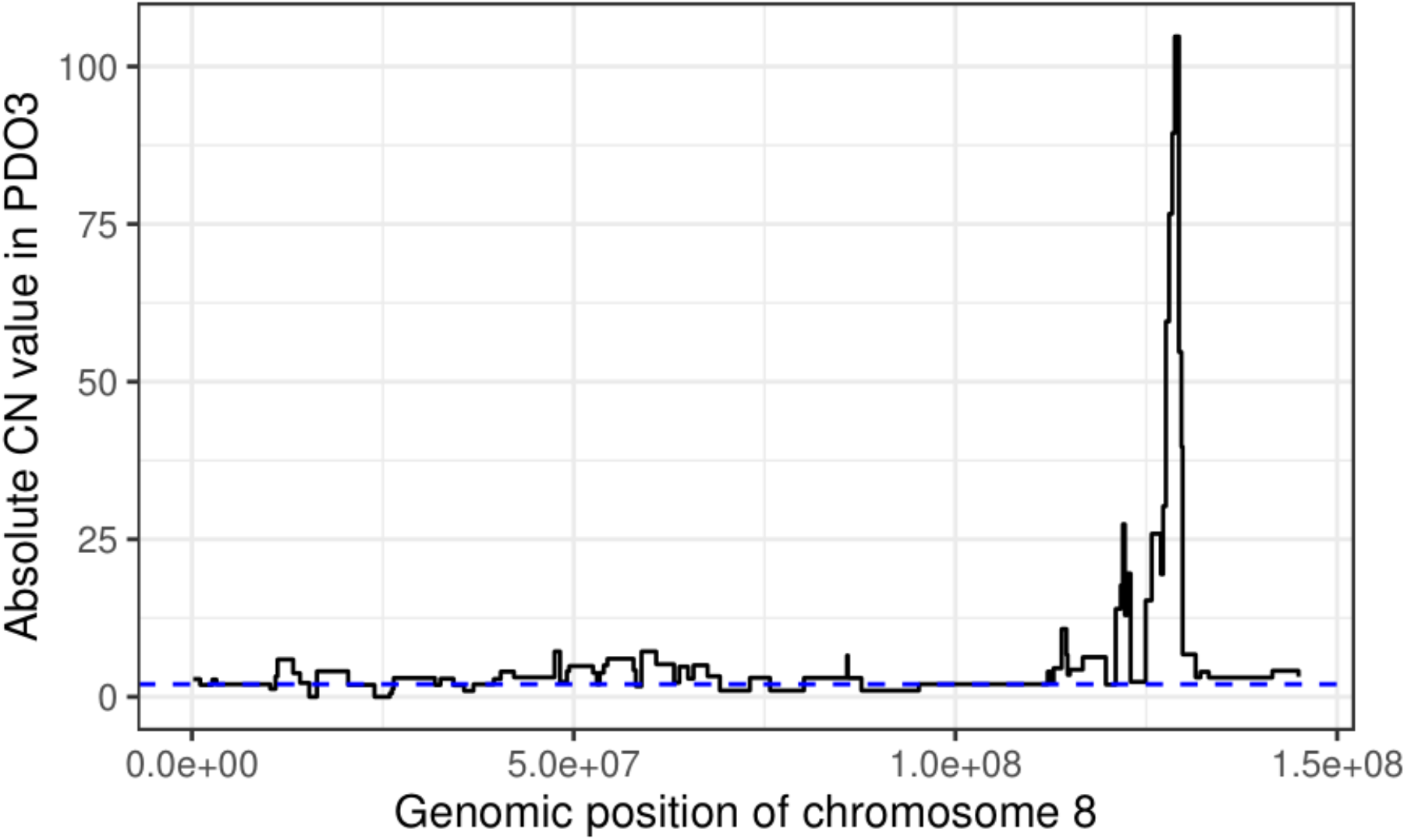
Chromosome 8 chromothripsis in PDO3. Step plot of the absolute copy number from sWGS for PDO3, along chromosome 8. The extreme values towards the end of the chromosome indicate a chromothriptic event. Diploid state is represented by the blue dotted line.

**Supplementary Fig. 12.**
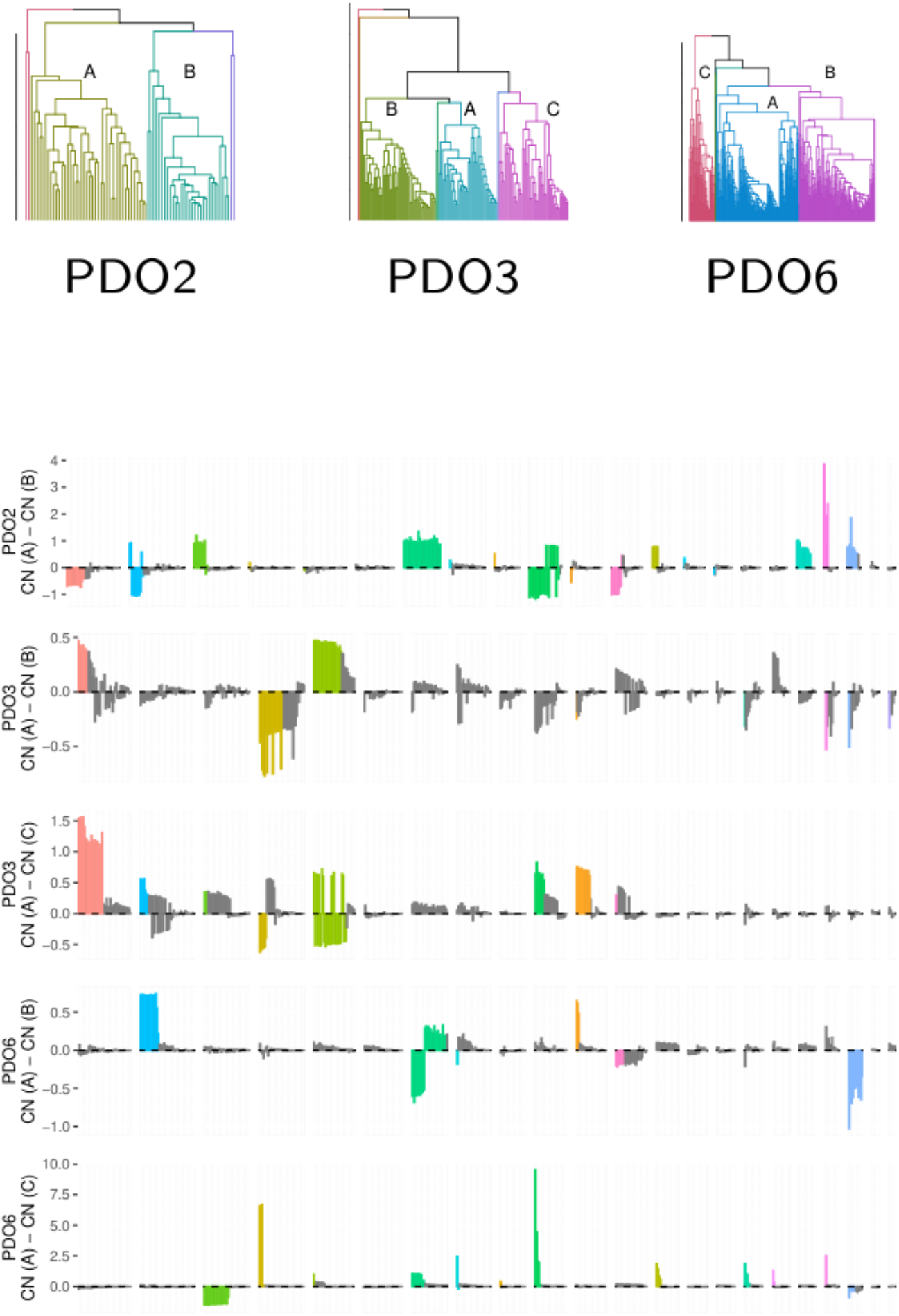
Major clades of cells in three organoids determined by copy number alterations from single cell DNA-seq. PDO2 contains two major clades of 42 and 30 cells. These clades are characterised by changes in chromosomes 7 (triplicated in the first clade and diploid in the second, in accordance with the copy number of roughly 2.75 from bulk sequencing) and chromosome 10 (showing the opposite trend). Few subclonal losses are observed, except for those in chromosome 13p. In PDO3, the two clades are comprised of 48 and 92 cells, with the second clade containing two subclades of 40 and 52 cells. The largest differences are in chromosomes 1, 4 and 5, 10 and 11. There is an LOH region at the start of chromosome 5 in the second clade, in all other chromosomes there are further copy number gains in regions where the first clade already shows amplifications. The two subclades contain differences in chromosomes 1, 4 and 5. PDO6 contains two clades of 49 and 303 cells (the second clade is split in subclades of 158 and 145 cells). The differences in the first split are in chromosomes 3, 4, 6, and 10, in which the extent of amplifications varies, and in chromosome 13, which presents large regions of single copy loss in the second clade. The greatest difference between the subclades is a large triplication in the first subclade of chromosome 2.

**Supplementary Fig. 13.**
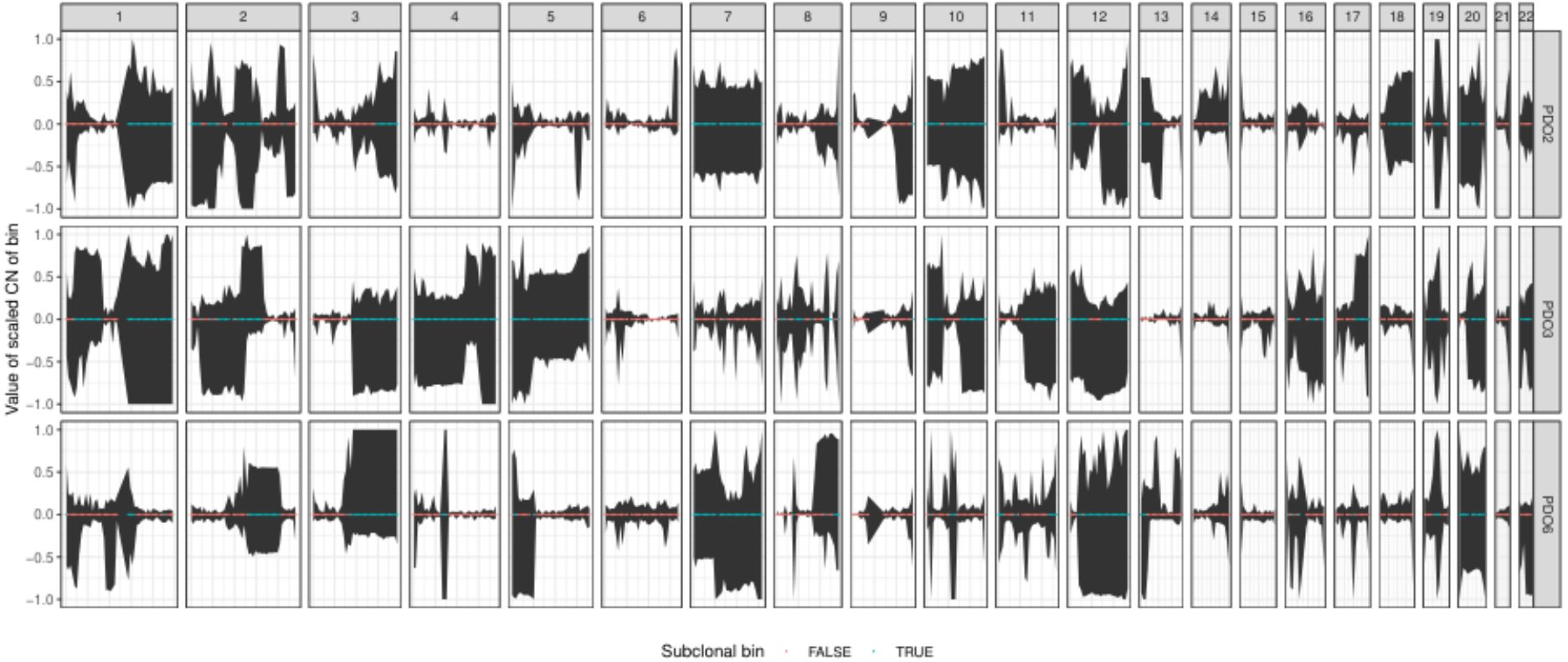
95% confidence intervals of the centered number across single cells from organoids along the genome. The colour of the dots indicates whether the CN value is shared between cells, and therefore clonal (in red), or whether there is subclonal heterogeneity in the bin (in blue) according to a Chi-squared test of the variance in copy number among cells and correcting for ploidy.

## Supplementary Tables

**Supplementary Table 1.**
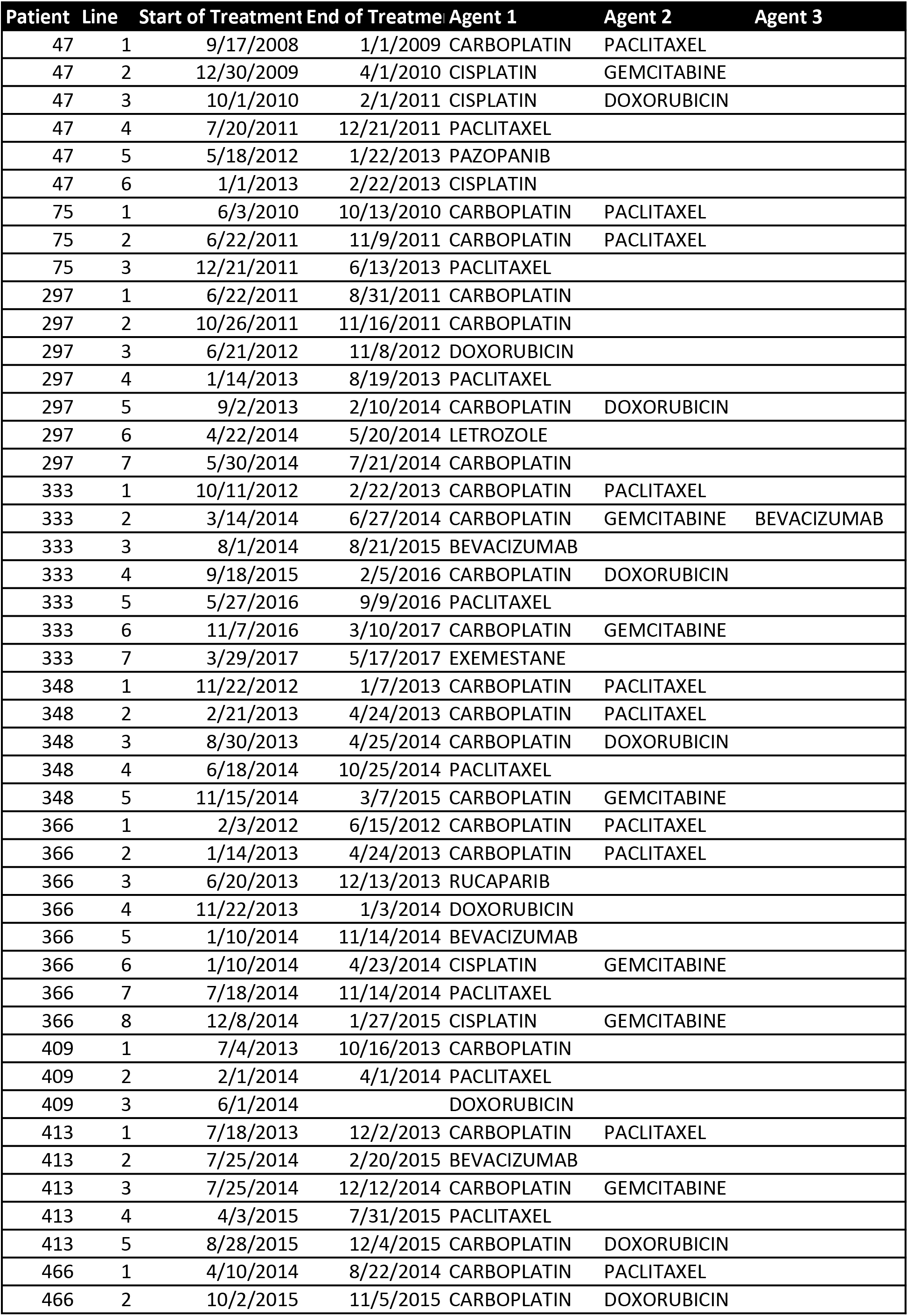
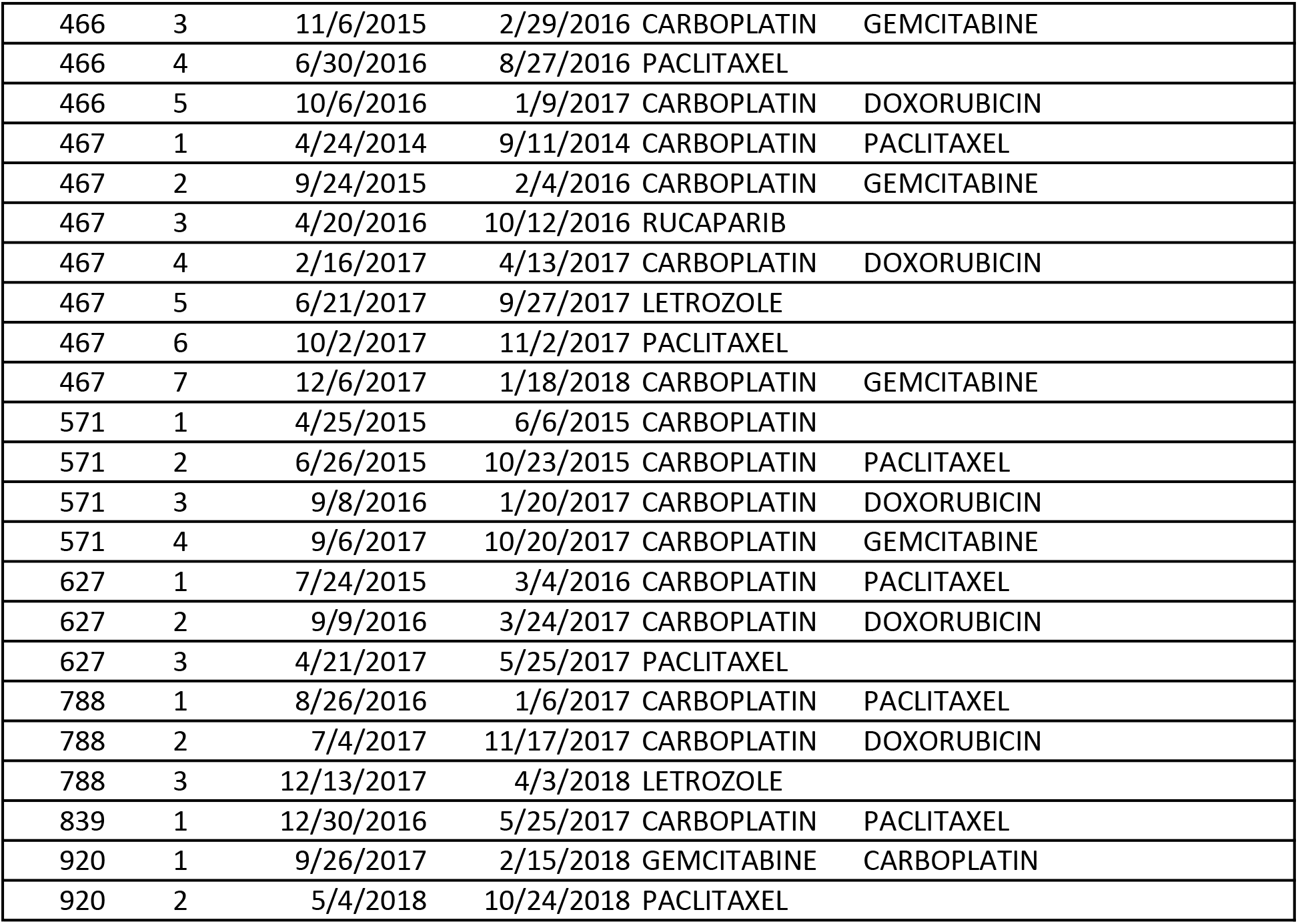
Summary of patient chemotherapy treatment

| Patient | Line | Start of Treatment | End of Treatment | Agent 1 | Agent 2 | Agent 3 |
| --- | --- | --- | --- | --- | --- | --- |
| 47 | 1 | 9/17/2008 | 1/1/2009 | CARBOPLATIN | PACLITAXEL |  |
| 47 | 2 | 12/30/2009 | 4/1/2010 | CISPLATIN | GEMCITABINE |  |
| 47 | 3 | 10/1/2010 | 2/1/2011 | CISPLATIN | DOXORUBICIN |  |
| 47 | 4 | 7/20/2011 | 12/21/2011 | PACLITAXEL |  |  |
| 47 | 5 | 5/18/2012 | 1/22/2013 | PAZOPANIB |  |  |
| 47 | 6 | 1/1/2013 | 2/22/2013 | CISPLATIN |  |  |
| 75 | 1 | 6/3/2010 | 10/13/2010 | CARBOPLATIN | PACLITAXEL |  |
| 75 | 2 | 6/22/2011 | 11/9/2011 | CARBOPLATIN | PACLITAXEL |  |
| 75 | 3 | 12/21/2011 | 6/13/2013 | PACLITAXEL |  |  |
| 297 | 1 | 6/22/2011 | 8/31/2011 | CARBOPLATIN |  |  |
| 297 | 2 | 10/26/2011 | 11/16/2011 | CARBOPLATIN |  |  |
| 297 | 3 | 6/21/2012 | 11/8/2012 | DOXORUBICIN |  |  |
| 297 | 4 | 1/14/2013 | 8/19/2013 | PACLITAXEL |  |  |
| 297 | 5 | 9/2/2013 | 2/10/2014 | CARBOPLATIN | DOXORUBICIN |  |
| 297 | 6 | 4/22/2014 | 5/20/2014 | LETROZOLE |  |  |
| 297 | 7 | 5/30/2014 | 7/21/2014 | CARBOPLATIN |  |  |
| 333 | 1 | 10/11/2012 | 2/22/2013 | CARBOPLATIN | PACLITAXEL |  |
| 333 | 2 | 3/14/2014 | 6/27/2014 | CARBOPLATIN | GEMCITABINE | BEVACIZUMAB |
| 333 | 3 | 8/1/2014 | 8/21/2015 | BEVACIZUMAB |  |  |
| 333 | 4 | 9/18/2015 | 2/5/2016 | CARBOPLATIN | DOXORUBICIN |  |
| 333 | 5 | 5/27/2016 | 9/9/2016 | PACLITAXEL |  |  |
| 333 | 6 | 11/7/2016 | 3/10/2017 | CARBOPLATIN | GEMCITABINE |  |
| 333 | 7 | 3/29/2017 | 5/17/2017 | EXEMESTANE |  |  |
| 348 | 1 | 11/22/2012 | 1/7/2013 | CARBOPLATIN | PACLITAXEL |  |
| 348 | 2 | 2/21/2013 | 4/24/2013 | CARBOPLATIN | PACLITAXEL |  |
| 348 | 3 | 8/30/2013 | 4/25/2014 | CARBOPLATIN | DOXORUBICIN |  |
| 348 | 4 | 6/18/2014 | 10/25/2014 | PACLITAXEL |  |  |
| 348 | 5 | 11/15/2014 | 3/7/2015 | CARBOPLATIN | GEMCITABINE |  |
| 366 | 1 | 2/3/2012 | 6/15/2012 | CARBOPLATIN | PACLITAXEL |  |
| 366 | 2 | 1/14/2013 | 4/24/2013 | CARBOPLATIN | PACLITAXEL |  |
| 366 | 3 | 6/20/2013 | 12/13/2013 | RUCAPARIB |  |  |
| 366 | 4 | 11/22/2013 | 1/3/2014 | DOXORUBICIN |  |  |
| 366 | 5 | 1/10/2014 | 11/14/2014 | BEVACIZUMAB |  |  |
| 366 | 6 | 1/10/2014 | 4/23/2014 | CISPLATIN | GEMCITABINE |  |
| 366 | 7 | 7/18/2014 | 11/14/2014 | PACLITAXEL |  |  |
| 366 | 8 | 12/8/2014 | 1/27/2015 | CISPLATIN | GEMCITABINE |  |
| 409 | 1 | 7/4/2013 | 10/16/2013 | CARBOPLATIN |  |  |
| 409 | 2 | 2/1/2014 | 4/1/2014 | PACLITAXEL |  |  |
| 409 | 3 | 6/1/2014 |  | DOXORUBICIN |  |  |
| 413 | 1 | 7/18/2013 | 12/2/2013 | CARBOPLATIN | PACLITAXEL |  |
| 413 | 2 | 7/25/2014 | 2/20/2015 | BEVACIZUMAB |  |  |
| 413 | 3 | 7/25/2014 | 12/12/2014 | CARBOPLATIN | GEMCITABINE |  |
| 413 | 4 | 4/3/2015 | 7/31/2015 | PACLITAXEL |  |  |
| 413 | 5 | 8/28/2015 | 12/4/2015 | CARBOPLATIN | DOXORUBICIN |  |
| 466 | 1 | 4/10/2014 | 8/22/2014 | CARBOPLATIN | PACLITAXEL |  |
| 466 | 2 | 10/2/2015 | 11/5/2015 | CARBOPLATIN | DOXORUBICIN |  |

Supplementary Table 1 | Summary of patient chemotherapy treatment
|  |  |  |  |  |  |
| --- | --- | --- | --- | --- | --- |
| 466 | 3 | 11/6/2015 | 2/29/2016 | CARBOPLATIN | GEMCITABINE |
| 466 | 4 | 6/30/2016 | 8/27/2016 | PACLITAXEL |  |
| 466 | 5 | 10/6/2016 | 1/9/2017 | CARBOPLATIN | DOXORUBICIN |
| 467 | 1 | 4/24/2014 | 9/11/2014 | CARBOPLATIN | PACLITAXEL |
| 467 | 2 | 9/24/2015 | 2/4/2016 | CARBOPLATIN | GEMCITABINE |
| 467 | 3 | 4/20/2016 | 10/12/2016 | RUCAPARIB |  |
| 467 | 4 | 2/16/2017 | 4/13/2017 | CARBOPLATIN | DOXORUBICIN |
| 467 | 5 | 6/21/2017 | 9/27/2017 | LETROZOLE |  |
| 467 | 6 | 10/2/2017 | 11/2/2017 | PACLITAXEL |  |
| 467 | 7 | 12/6/2017 | 1/18/2018 | CARBOPLATIN | GEMCITABINE |
| 571 | 1 | 4/25/2015 | 6/6/2015 | CARBOPLATIN |  |
| 571 | 2 | 6/26/2015 | 10/23/2015 | CARBOPLATIN | PACLITAXEL |
| 571 | 3 | 9/8/2016 | 1/20/2017 | CARBOPLATIN | DOXORUBICIN |
| 571 | 4 | 9/6/2017 | 10/20/2017 | CARBOPLATIN | GEMCITABINE |
| 627 | 1 | 7/24/2015 | 3/4/2016 | CARBOPLATIN | PACLITAXEL |
| 627 | 2 | 9/9/2016 | 3/24/2017 | CARBOPLATIN | DOXORUBICIN |
| 627 | 3 | 4/21/2017 | 5/25/2017 | PACLITAXEL |  |
| 788 | 1 | 8/26/2016 | 1/6/2017 | CARBOPLATIN | PACLITAXEL |
| 788 | 2 | 7/4/2017 | 11/17/2017 | CARBOPLATIN | DOXORUBICIN |
| 788 | 3 | 12/13/2017 | 4/3/2018 | LETROZOLE |  |
| 839 | 1 | 12/30/2016 | 5/25/2017 | CARBOPLATIN | PACLITAXEL |
| 920 | 1 | 9/26/2017 | 2/15/2018 | GEMCITABINE | CARBOPLATIN |
| 920 | 2 | 5/4/2018 | 10/24/2018 | PACLITAXEL |  |

**Supplementary Table 2.** Cell lines derived from PDOs

| Patient-derived organoid | Cell line | Patient ID |
| --- | --- | --- |
| PDO3 | CIOV4 | 466 |
| PDO2 | CIOV5 | 75 |
| PDO7 | CIOV6 | 366 |
| PDO1 | CIOV7 | 920 |

**Supplementary Table 3.**
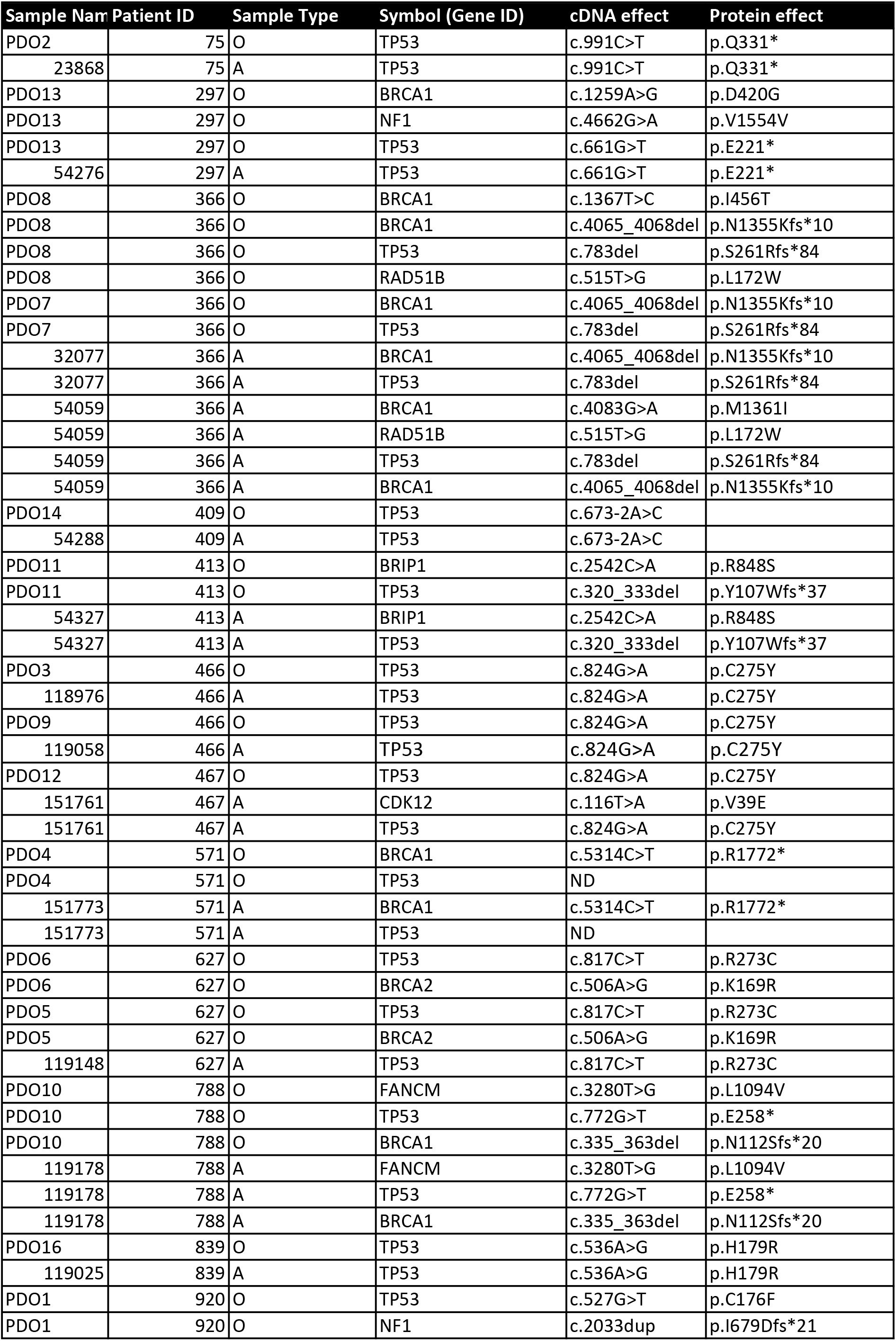
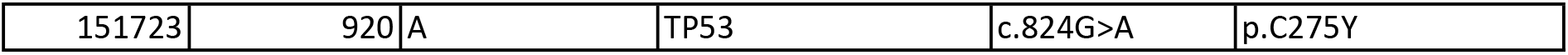
Mutation analysis of patient and PDO samples

**Supplementary Table 4.**
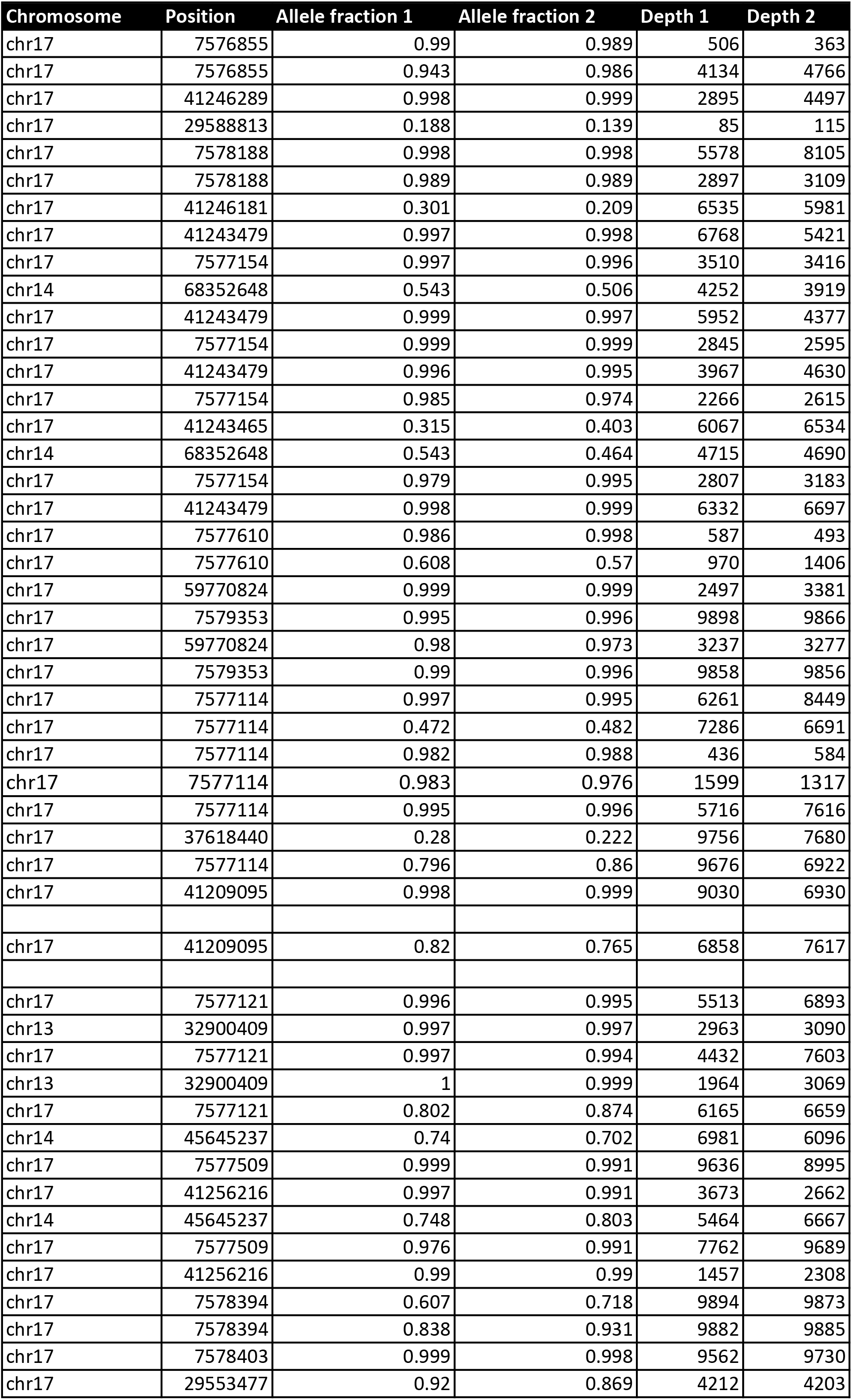
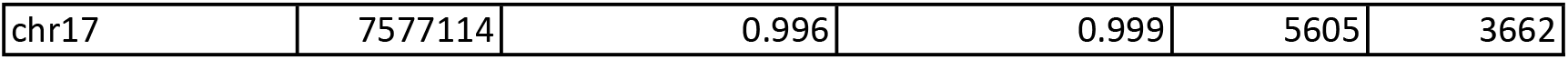
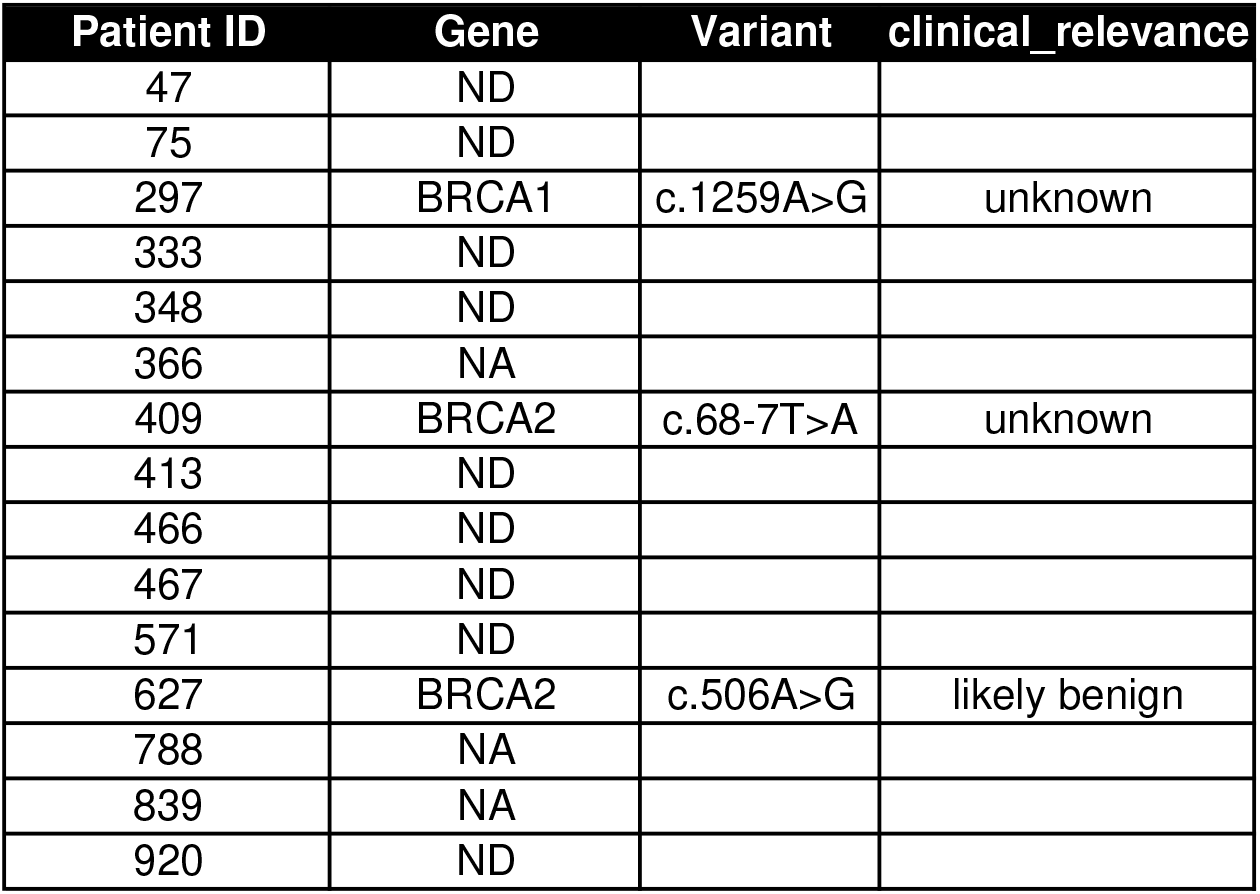
Patient germline BRCA mutation status

